# Germline protein, Cup, non-cell autonomously limits migratory cell fate in *Drosophila* oogenesis

**DOI:** 10.1101/2022.07.11.499544

**Authors:** Banhisikha Saha, Sayan Acharjee, Gaurab Ghosh, Purbasa Dasgupta, Mohit Prasad

## Abstract

Attaining migratory fate from a stationary cell population is complex and indispensable both for the multicellular organism development as well for the pathological condition like tumor metastasis. Though widely prevalent in the metazoans, the molecular understanding of this phenomenon remains elusive. Specification of migratory border cells from the follicular epithelium during *Drosophila* oogenesis has emerged as one of the excellent model systems to study how motile cell are specified. JAK-STAT activation in 6-10 anterior most follicle cells of the *Drosophila* egg chamber transforms them to a migratory cluster called the border cells. We show that a nurse cell protein, Cup, non-cell autonomously restricts the domain of JAK-STAT activation in the anterior follicle cells. Further examination suggests that Cup functions through Rab11GTPase to regulate Delta trafficking in the nurse cells potentiating Notch activation in the anterior follicle cells. Since Notch activity in the follicle cells modulates the JAK-STAT, any perturbation in Notch activation affects the border cell fate. Altogether, we propose that germline Cup affects the border cell fate through appropriate activation of Notch and JAK-STAT signaling in the follicle cells.

## Introduction

Acquisition of migratory fate from a stationary epithelium not only plays an important role in aiding normal metazoan development but is also linked to various pathological conditions including tumor cell metastasis. (Ciruna and Rossant 2001; Jiang et al. 2013; Perrimon, Pitsouli, and Shilo 2012). Unfortunately, unwarranted specification of migratory cells from solid tumours is one of the leading causes of fatality associated with cancer metastasis (Friedl and Gilmour 2009; Naora and Montell 2005; Rørth 2009; Thiery et al. 2009). Although cells employ diverse mechanisms to acquire migratory fates, broadly they can be classified either under an autonomous or the regulative mode of specification (Davidson, Cameron, and Ransick 1998; Edlund and Jessell 1999). Unlike autonomous mode, regulative communication dominates cell fate specification because of plethora of diverse cell-cell interactions possible in the metazoans. Since the transition of epithelial fate to mesenchymal fate is a prerequisite for growth, development, and survival, there is constant attempt to understand how migratory group of cells is delineated from their static progenitors in the multicellular organisms.

Border cells (BCs) in *Drosophila* oogenesis has emerged as an excellent genetic model system for studying how motile cells are specified from a stationary epithelium (Denise J Montell 2001). *Drosophila* oogenesis is a synchronised developmental process consisting of 14 stages of interconnected oval egg chambers (Bastock and St Johnston 2011; Spradling 1993). Each egg chamber harbours 16 central germline cells, of which 1 takes the oocyte fate, while the rest acquires nurse cell identity that nourishes the growing oocyte (Horne-Badovinac and Bilder 2005; Huynh and St Johnston 2004; Denise J. Montell 2003). Enveloping the germline cells is a single layer of approximately 750 follicle epithelial cells. A pair of specialized follicle cells called the polar cells mark each end of the egg chamber (Ruohola et al. 1991). The polar cells secrete cytokine,Unpaired (Upd) that activates JAK-STAT pathway and aids in specifying migratory fate to a select group of 4-6 anterior follicle cells (AFCs) (McGregor, Xi, and Harrison 2002; Silver and Montell 2001). This migratory group of AFCs that undergo partial Epithelial to Mesenchymal fate transition and initiate posterior movement towards the oocyte are the BCs (Denise J. Montell 2003). Activation of CEBP transcription factor, Slow border cells (Slbo), by JAK-STAT marks the fate of BCs (Beccari, Teixeira, and Rørth 2002; D J Montell, Rorth, and Spradling 1992; Silver and Montell 2001). After the BCs are specified their posterior movement is guided through a gradient of growth factors (PVF1-Platelet Derived Growth Factor and Vascular Endothelial Growth Factor-related Factor 1and Egf-Epidermal growth factor) secreted from the oocyte (P Duchek and Rorth 2001; Peter Duchek et al. 2001; McDonald 2003). Once the cluster reaches the oocyte boundary, it aids in the formation of a channel in the micropyle, which permits the sperm entry during fertilization (D J Montell, Rorth, and Spradling 1992). Any defect in BC specification/ cluster formation or their efficient movement, impedes micropyle function, thus rendering the egg sterile.

The JAK-STAT signaling in the AFCs is strictly modulated at multiple steps to recruit an optimum number of FCs to BCs fate (generally 4-6 cells). At the primary level, both the production and the distribution of Upd ligand are regulated to form a gradient across the anterior follicle cells. Yorkie, a component of the Hippo signaling pathway negatively regulates Upd production from the polar cells (Lin et al. 2014). On the other hand, the Glypicans, Dally, and Dally-like shape the distribution of Upd ligand, thus affecting STAT signaling and BC fate specification (Hayashi et al. 2012). Further within the AFCs, various intracellular components modulate STAT activity. Suppressor of Cytokine Signaling (SOCS36E) regulates ubiquitination of several components of the JAK-STAT pathway to limit STAT activation (Monahan and Starz-Gaiano 2015; Stec, Vidal, and Zeidler 2013). In addition, there are checkpoints at the transcriptional level too. In the follicle cells (FCs), antagonistic interactions between STAT and transcriptional repressor Apontic, restricts the domain of STAT activation, thereby limiting BC fate specification (Starz-Gaiano et al. 2008). A recent study shows that Insulin signaling limits BC fate by stabilising the negative regulator SOCS36E in the AFCs (Kang et al. 2018). Thus, the JAK-STAT pathway is regulated through multiple ways in the somatic FCs to limit BC fate during *Drosophila* oogenesis. Since interaction between germline nurse cells and somatic FCs is critical for oogenesis progression and polar cell fate specification, we were curious to examine if the germline cells have any direct role in modulation of BC specification (Assa-Kunik et al. 2007).

In this study, we report a novel role of germline nurse cells in BC fate specification. Specifically, our data suggest that Cup protein, which expresses in the germline, non-cell autonomously modulates Notch signaling in the FCs. As Notch and JAK-STAT signaling are antagonistic, Cup mutants exhibit excess BC fate due to elevated STAT in the AFCs. Further, we demonstrate that Cup mutants exhibit disturbed actin cytoskeleton network and enrichment of Delta puncta in the nurse cell cytoplasm. Employing classical genetics and tissue immunohistochemistry in various genetic backgrounds, we propose that Cup maintains the germline cytoskeletal integrity and modulates Delta trafficking in the nurse cells. As we observe rescue in the BC numbers when constitutively active Rab11GTPase is overexpressed in the Cup mutant germline, we propose that recycling Delta in the germline nurse cells is critical for Notch activation in the AFCs of vitellogenic egg chambers. Notch stimulation in the AFCs modulates STAT activity, thus controlling the number of AFCs that acquire BC fate.

## Results

### Germline function of Cup affects the size of somatic BC cluster

BC specification and migration is one of the critical factors that determine the fertility of the female flies. There are several autocrine, and paracrine factors associated with AFC that mediate specification of BCs. Though signals from the nurse cells regulate the polar and stalk cell fate in previtellogenic egg chambers (stage 1-2), it is not clear if the germline can directly impact the specification of BCs (López-Schier and St. Johnston, 2001a). Hence, we enquired if the germline nurse cells participate in the specification of somatic BCs during early vitellogenesis.

To address this question, we shortlisted 14 genes which are known to be expressed in the nurse cells and their mutants are reported to be female sterile. (Table S1). Among the 14 genes, we examined the status of BC fate specification in 3 mutant lines that were known to be homozygous viable. We evaluated the size of BC clusters in the respective homozygous mutant egg chambers with the premise that any alteration in BC fate would have a direct bearing on the cluster size. We measured the size of the BC cluster for each of the three homozygous mutant lines and found that mutation in the Cup gene (*cup^01355^*) resulted in the largest BC cluster (3073.40±127.59µm^3^ SEM, n=32 clusters) compared to the WT (1373.33±54.86 µm^3^ SEM, n=31 clusters) (Fig. 1A-C). The description of the cluster size of other mutant lines is provided in Fig. S1A-D.

**Figure 1:**
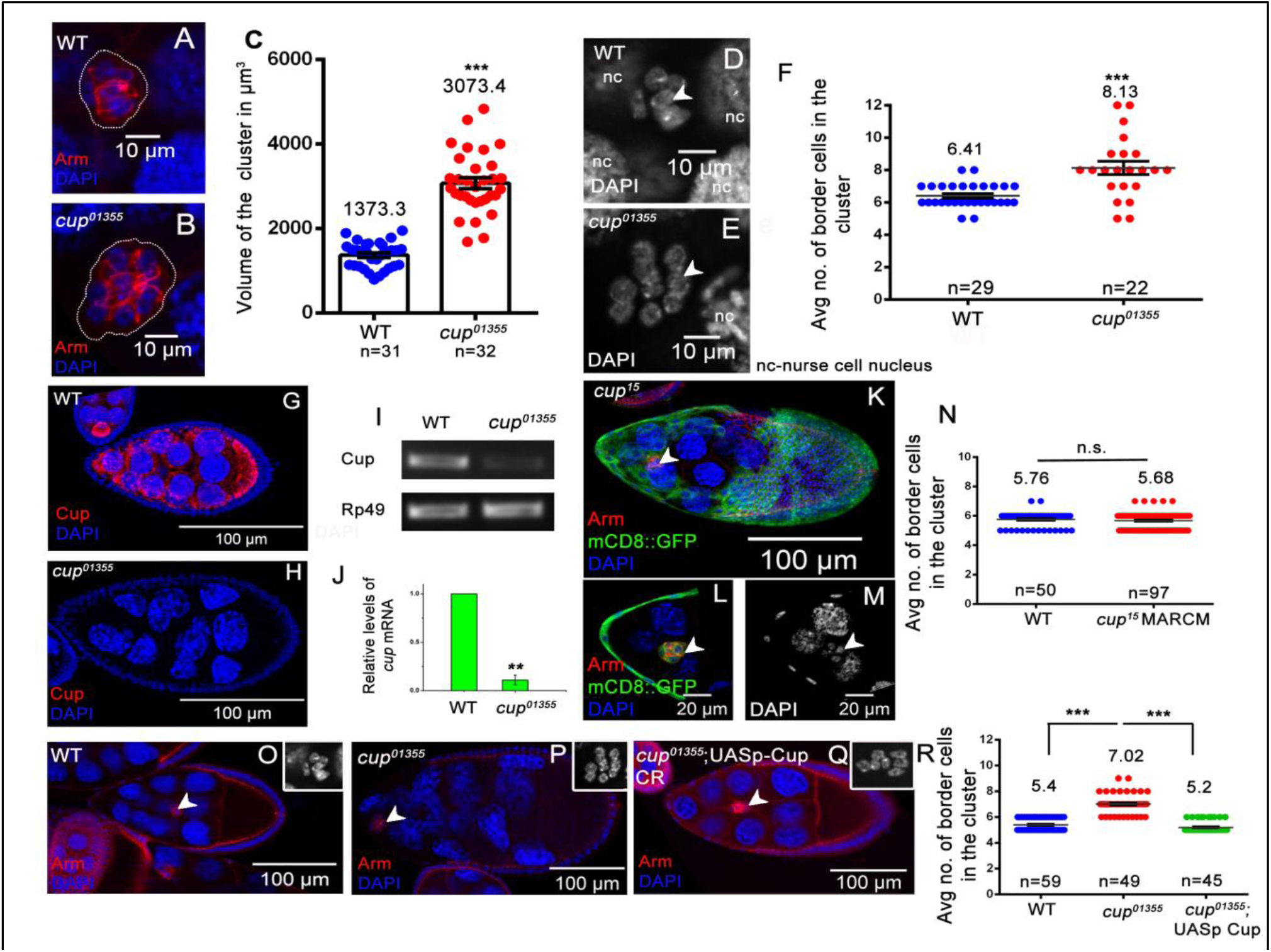
Cup functions in the germline cells to affect the size of the BC cluster. (**A-C)** *cup^01355^*egg chambers exhibit increased border cell cluster size, Armadillo (red), DAPI (blue) and cell count (**D-F**), DAPI (grey), compared to wild type. White dotted line marks BC cluster. (**G-H**) *cup^01355^* egg chambers lack Cup expression, Cup (red), DAPI (blue) and exhibit reduced *cup* transcripts, normalised to *rp49* **(I-J)** compared to wild type. (**K-N**) AFC clones mutant for *cup^15^* marked by GFP(green), Armadillo (red), DAPI (blue and grey) does not alter number of BCs (white arrow) compared to control. (**O-R**) Increased BC number (white arrow) is rescued by UASp-Cup CR (Coding Region), driven by *nos.NGT*-Gal4 in *cup^01355^* egg chambers, Armadillo (red), DAPI (blue, inset grey).

*cup^01355^* is a hypomorphic allele of the Cup gene that regulates the translation and stability of several maternal mRNAs including that of *oskar* and *nanos* during *Drosophila* oogenesis(Broyer, Monfort, and Wilhelm 2017; Nelson, Leidal, and Smibert 2004; Wilhelm et al. 2003) This allele has a *P-lacZ* insertion in the untranslated region of the first exon of the Cup gene and belongs to the least severe class of alleles, where the phenotype manifests during post vitellogenic stages of *Drosophila* oogenesis (Keyes and Spradling, 1997). This mutant was ideal for our analysis as all other approaches to down regulate Cup function in the germline stalled the egg chambers in early stages of oogenesis.

To test if the larger clusters observed in *cup^01355^* homozygotes were indeed due to the altered number of BCs, we stained the egg chambers with DAPI to quantify the number of BCs. Consistent with our expectation, we observed that the number of BCs in *cup^01355^* mutant egg chambers was higher (8.13±0.4 SEM, n=22) compared to WT (6.41±0.13 SEM, n=29) (Fig. 1D-F). This suggested that Cup probably modulates the number of border cells in the migrating cluster in developing egg chambers.

Since *cup^01355^* mutants egg chambers include both germline and somatic FCs, we were curious to know in which cells was the Cup protein indeed functioning that was modulating border cell fate in AFCs. We immunostained the egg chambers with an anti-Cup antibody and observed that Cup is highly expressed in the cytoplasm of the germline cells both in the early and late stages of oogenesis. Consistent with previous published reports, we failed to detect any Cup protein in the somatic FCs (Keyes and Spradling, 1997). Since *cup^01355^* is a hypomorphic allele, we examined the levels of Cup transcript and protein in *cup^01355^* ovaries. Though we observed reduced levels of *cup* transcript (1/10th of wild type), we failed to detect any Cup protein in the *cup^01355^* egg chambers (Fig. 1G-J). The expression analysis of Cup gene product in WT and *cup^01355^* egg chambers suggested that Cup is primarily a germline protein and probably non-cell autonomously affecting BC fate in the FCs. To further confirm this, we employed Mosaic Analysis with a Repressible Cell Marker (MARCM) technique to generate homozygous mutant Cup FCs using a stronger allele of Cup (*cup^15^*) and examined the status of BC fate specification (Lee and Luo, 2001). *cup^15^* is an EMS allele, and mutant ovaries are known to exhibit a negligible amount of Cup protein as compared to WT (Keyes and Spradling, 1997). As expected, we didn’t observe any significant difference in BC numbers specified in *cup^15^* mutant AFCs (5.68±0.06 SEM, n=97) compared to WT AFCs (5.76±0.07 SEM, n=50) (Fig. 1K-N). Further, we downregulated Cup function by generating flip-out clones expressing *cup* RNAi spanning the entire AFCs with at least 4 BC clones and quantified the number of BCs (Menon et al., 2015). Similar to our MARCM analysis, we did not observe any alteration in BC specification due to *cup* RNAi overexpression (5.4±0.08 SEM, n=50) compared to control (5.45±0.08 SEM, n=51) (Fig. S1E-H). We also downregulated Cup function in BC precursor FCs by expressing *cup* RNAi employing *c306-*GAL4 driver and observed no difference in BC numbers (5.25±0.05 SEM, n=76) compared to in observed control egg chambers (5.26±0.05 SEM, n=75) (Fig. S1I-K). Finally, to validate that the increased BC number is indeed due to the absence of Cup in the nurse cells, we restored Cup expression by expressing the Cup-coding region (Cup-CR) in *cup^01355^* nurse cells using *nos.NGT* GAL4. Upon reconstitution of Cup-CR in *cup^01355^* nurse cells, the BC number was significantly restored close to that of the WT(*cup^01355^*-7.02±0.11SEM, rescue-5.2±0.06 SEM, wild type-5.4±0.06 SEM, n≥45 egg chambers) (Fig. 1O-R).

Altogether our results above suggest that we have identified a nurse cell protein, Cup, which non-cell autonomously modulates the size of BC cluster specified from the overlying somatic FCs.

### Cup controls BC fate by negatively regulating the JAK-STAT pathway

Since *cup^01355^* mutant egg chambers exhibit more nuclei in the migrating cluster, we investigated if the extra cells were indeed BCs. To check this, we stained the egg chambers with the Slbo antibody, which conspicuously marks the BCs. We observed significantly higher number of Slbo positive cells in the cluster (7.04±0.19 SEM, n=23) in *cup^01355^* egg compared to the WT (5.27±0.11SEM, n=22) (Fig. 2A-C). This suggests that Cup mutation results in aberrant specification of BCs from the follicular epithelium. To rule out the possibility that an increase in BC numbers is due to altered endoreplication of FCs, we compared the expression pattern of endoreplication markers Cut and phospho Histone 3 (pH3) between the WT and *cup^01355^* egg chambers (López-Schier and St. Johnston, 2001; Sun, 2005). We examined 170 egg chambers each of WT, and *cup^01355^* and observed no difference in staining pattern for Cut (Fig. S2A-H). We didn’t observe any pH3 positive cells in stage 8, or later egg chambers in 168 samples analyzed each for WT and *cup^01355^* (Fig. S2I-L”). As the expression pattern of both Cut and pH3 was similar in both the WT and the *cup^01355^* egg chambers, we excluded the possibility of altered endoreplication being the cause of excessive BC fate in the Cup mutants.

**Figure 2.**
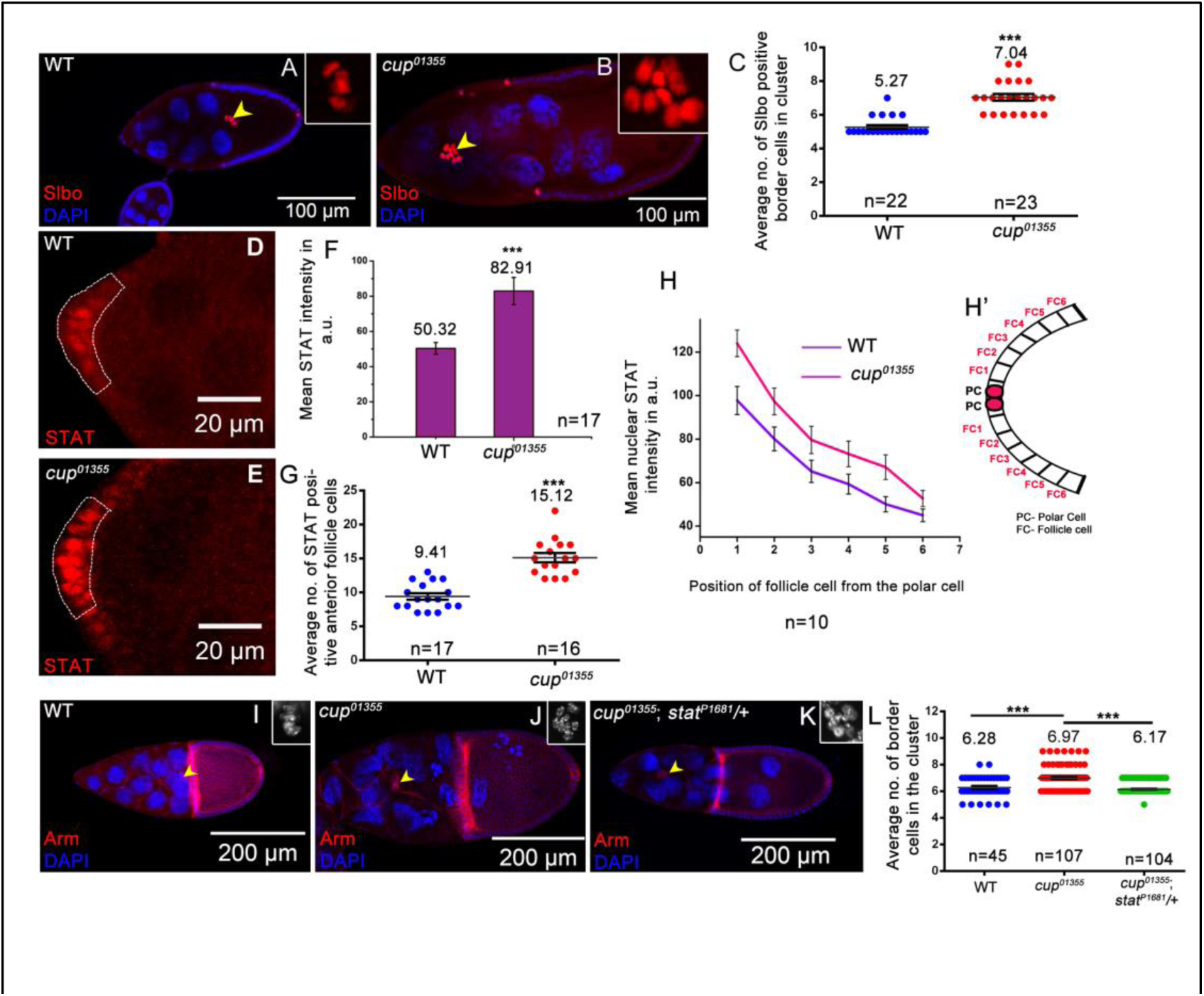
Cup controls BC fate by negatively regulating the JAK-STAT pathway. (**A-C**) *cup^01355^* egg chambersexhibit increased Slbo positive cells (yellow arrow), Slbo (red), DAPI (blue), compared to wild type. (**D-G**) *C* egg chambers exhibit higher STAT levels and more STAT positive cells in anterior end of egg chamber (dotted area), STAT (red) as compared to wild type. (**H-H’**) STAT level is higher in 6^th^ FC from polar cell in *cup^01355^*egg chambers as compared to control. (**I-L**) Higher BC numbers in *cup^01355^* is rescued when *stat^p1681^/+* background is introduced (yellow arrows), Armadillo (red), DAPI (blue, inset grey).

Since JAK-STAT signaling activates Slbo expression in the AFCs, we next examined if the increase in the number of BCs was linked to enhanced STAT function (Beccari et al., 2002; Silver and Montell, 2001). Nuclear STAT is used as a molecular reporter for assessing the status of JAK-STAT signaling (Darnell et al., 1994). To measure STAT activity, we quantified nuclear STAT in WT and *cup^01355^* mutant AFCs. Unlike WT (50.32±3.36 SEM, n=17), we observed higher levels (1.64-fold) of STAT in *cup^01355^* mutant FCs (82.91±7.76 SEM, n=17 egg chambers) (Fig. 2D-F). Also, we observed that the number of AFCs exhibiting distinct nuclear STAT in *cup^01355^* egg chambers (15.12±0.67 SEM, n=16) was higher compared to WT (9.41±0.47 SEM, n=17) (Fig. 2G). In addition, we observed conspicuous nuclear STAT staining extending as far as 6th FC from the polar cell in *cup^01355^* egg chambers compared to 3 cells observed in the control (Fig. 2H-H’). These results suggest that both the levels and domain of STAT activation is enhanced in *cup^01355^* egg chambers. We then investigated if the elevated STAT was indeed responsible for excess BCs observed in the *cup^01355^* egg chambers. To test, this we compared BC numbers in *cup^01355^* egg chambers in WT and STAT heterozygous background (*stat^P1681^*/+). We observed BC number in *the cup^01355^* cluster was reduced in STAT heterozygous background than the Wild type (*cup^01355^*-6.97±0.077 SEM, *cup^01355^; stat^P1681^ /*+-6.17±0.03 SEM, wild type-6.28±0.11 SEM, n≥45 egg chambers) suggesting that elevated STAT is responsible for excess BC fate in the *cup^01355^* egg chambers (Fig. 2I-L). Given that higher STAT levels resulted excessive BC fate, we were eager to check as to how STAT function is elevated in *cup^01355^* FCs.

### Loss of Cup reduces Notch signaling, which leads to increased JAK-STAT activation

JAK-STAT signaling in the AFCs is positively regulated by the Upd ligand produced by the anterior polar cells (McGregor et al., 2002; Silver and Montell, 2001). Since we detected increased JAK-STAT signaling in the Cup mutant FCs, we examined if this was due to increase number of ligands producing polar cells. For this, we checked the pattern of Fasciclin III (FasIII), the lateral membrane protein that marks the junction between two polar cells (Ruohola et al., 1991). Like the wild type, we observed a single distinct junction of polar cells labelled by FasIII at the anterior and posterior ends in the *cup^01355^ egg* chambers suggesting normal number of polar cells (Fig. S3A-B). We also observed similar FasIII expression in early stages of oogenesis in both WT and *cup^01355^* egg chambers indicating that polar cell fate is unaffected in the *cup^01355^* hypomorphic background (Fig. S3C-J’). Given that the polar cell number is unaltered, we tested if enhanced JAK-STAT signaling was due to transcriptional upregulation of *the upd* gene itself. To examine this, we measured the expression of *the upd* reporter construct, *upd*-lacZ and observed no significant difference in the intensity of β-gal antibody staining from *upd*-lacZ between the WT (118.69±7.02 SEM, n=20) and the *cup^01355^* stage 8 egg chambers (124.73±11.03 SEM, n=20). This suggested against our premise that Upd gene expression was elevated in the *cup^01355^* egg chambers (Fig. S3K-M). A similar conclusion was made when β-gal antibody staining intensity was compared between the WT and the *cup^01355^* egg chambers in earlier stages of oogenesis (Fig. S3N-V). Altogether these results suggest that excess border cell fate observed in *cup^01355^* egg chambers is probably not due to alteration in the polar cell fate nor due to excessive transcriptional output from the Upd gene. This prompted us to explore the possibility of Cup modulating function of other JAK-STAT regulators that may, in turn, affect BC specification. One possible explanation for the upregulated STAT activity could be due to the downregulation of some of the negative regulators of the JAK-STAT signaling pathway in the *cup^01355^* egg chambers. There are several negative regulators of the JAK-STAT signaling including Protein tyrosine phosphatase 61F (Ptp61f), Brahma (Brm), Suppressor of Cytokine Signaling 36E (SOCS36E), and Notch (Assa-Kunik et al., 2007; Liu et al., 2010; Saadin and Starz-Gaiano, 2016). Among these molecules, we narrowed down to Notch primarily because of two reasons. The seat of Cup expression, the germline nurse cells is known to communicate with FCs via Notch signaling at several stages to permit egg chamber development. Secondly, the Notch signaling has been shown to act antagonistically to JAK-STAT signaling in a context-specific manner in the FCs (Assa-Kunik et al., 2007; López-Schier and St. Johnston, 2001a). Since we observed an upregulation of STAT function in the AFCs of *cup^01355^* egg chambers, we examined the level of Notch signaling in the FCs. For this we employed the Notch reporter construct where Notch Response Element (NRE) is tagged upstream of EGFP (NRE-EGFP). NRE comprises of binding sites for the Notch target, Suppressor of Hairless, and the transcriptional activator Grainy head (Zacharioudaki and Bray, 2014). Activation of Notch signaling leads to the binding of these transcriptional activators to the NRE sequence and results in GFP expression. We checked Notch activity in AFCs by measuring EGFP reporter expression (under NRE) in both WT and *cup^01355^* stage 8 egg chambers. Interestingly, we observed significantly lower levels of EGFP in the AFCs in *cup^01355^* egg chambers (6.29±0.38 SEM, n=20) compared to the WT (26.93±2.18 SEM, n=20) (Fig. 3A-C) suggesting that Notch signaling is severely compromised in the AFCs of *cup^01355^* egg chambers. To further support our observation with NRE-GFP, we examined the levels and distribution of Notch in the FC. It is known that ligand binding stimulates two sequential proteolytic cleavages in the Notch receptor generating a fragment with extracellular domain (NECD) and the other with intracellular domain (NICD) (Bray 2006). The distribution of NICD and NECD is routinely used to evaluate the status of Notch signaling. Ligand stimulation, promotes NECD and NICD internalization in the ligand producing cell and signal receiving cell respectively(Kopan and Ilagan 2009; Kovall et al. 2017; Nichols, Miyamoto, and Weinmaster 2007). We observed numerous NICD and NECD puncta in the wild type FC and Nurse cells respectively (Fig 3D-J). The presence of large number of NICD and NECD puncta suggests that Notch signaling is active in the wild type FCs. On the contrary, we observed very few internalized puncta of both NICD and NECD in the FC and the nurse cell of the Cup mutant egg chambers respectively supporting the fact the Notch signaling is downregulated (For NICD; WT-101±8.32 SEM n=10, *cup^01355^*-64.10±3.345 SEM n=10 and For NECD WT-271.9±24.18 SEM n=11, *cup^01355^*-103.4±15.30 SEM n=11. (Fig 3I,J).

**Figure 3.**
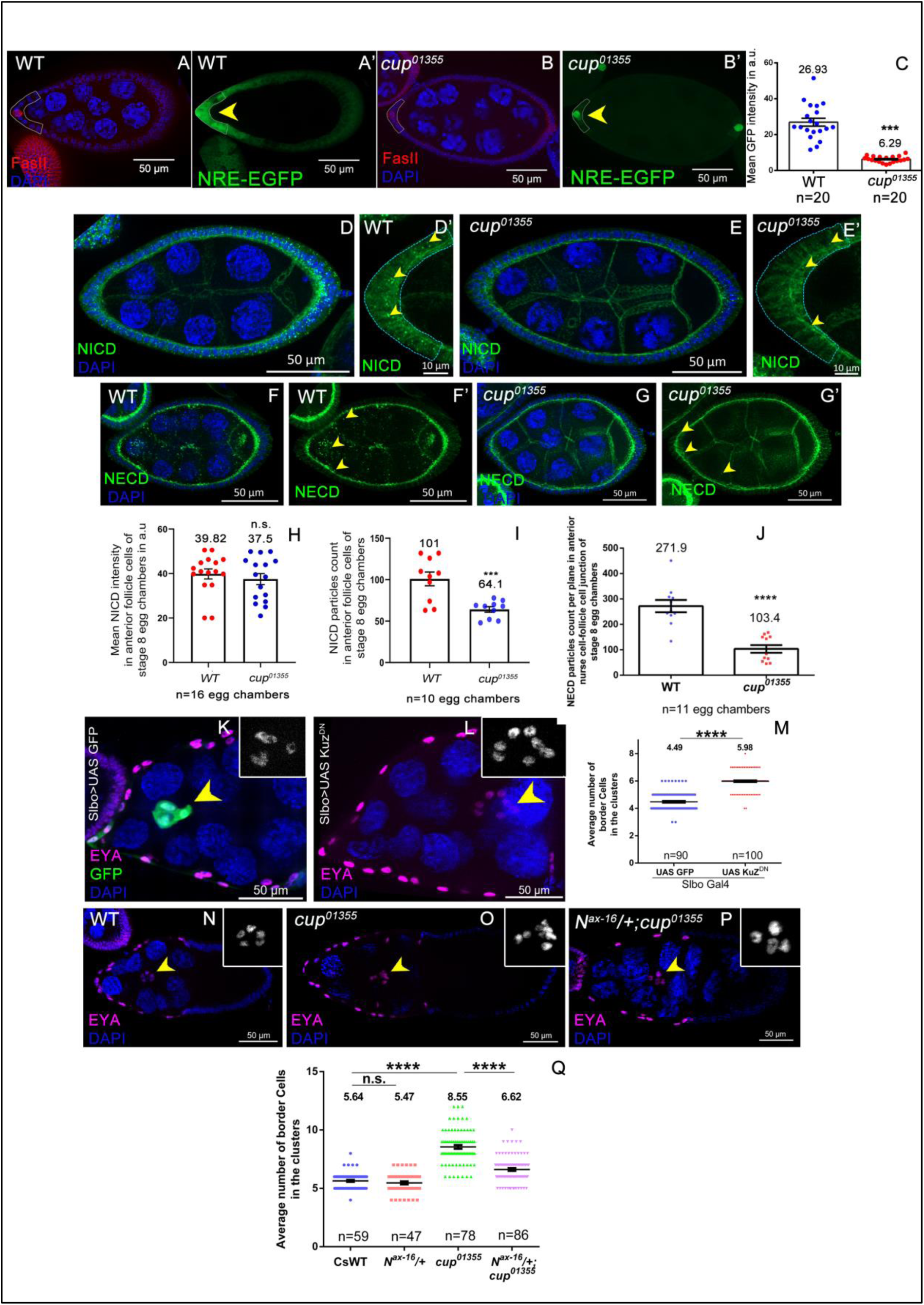
Loss of Cup reduces Notch signaling, which leads to increased JAK-STAT signaling. **(A-C)** Intensity of NRE -EGFP (in green) is significantly decreased in stage 8. *cup^012355^* egg chambers marked by dotted white line; Depleted FASII staining indicates stage 8 egg chamber DAPI indicated in Blue. **(D-E’)** NICD stained egg-chambers exhibit reduced number of puncta (yellow arrow) in follicle cells in *cup^012355^* with respect to WT. **(H-I)** Though mean intensity of NICD is not significantly different between *cup^012355^* and WT, the no of NICD puncta (yellow arrow) is reduced in anterior follicle cells in *cup^012355^* over WT. **(F-G’)** *cup^012355^* exhibits lesser number cytoplasmic NECD puncta (yellow arrow) with respect to Control. **(K-M)** Images of stage 9-10 egg chambers stained with Eya (in magenta) and DAPI(in blue)with indicated genotypes. Border cells (yellow arrow) express UAS mCD8GFP and UAS Kuz^DN^ by *Slbo-GAL4*. **(J)** Kuz^DN^ over-expression increases Border cells number over Control. **(N-Q)** Genetic interaction between Notch hyperactive allele (N^ax-16^) and *cup. N^Ax-16^*/+; *cup^012355^/ cup^012355^*gives partial rescue in BC number with respect to *cup^012355^/ cup^012355^* Yellow arrow border cells, Eya stain labels all FC and BC nuclei

Since Notch signaling was lower in *cup^01355^* egg chambers, we speculated if this condition caused excessive BC fate observed in the genotype of our interest. To test this possibility, we first down regulated Notch signaling in the AFCs. As Notch signaling is required in early oogenesis, conditional over expression of dominant negative Kuzbanian was carried out by *slbo*-GAL4 at 29°C to downregulate Notch signaling in stage 8 AFC just when BC fate is determined. Kuzbanian is a disintegrin metalloprotease that cleaves and releases the NICD fragement, thus activating Notch receptor (Lieber et al., 2002; Qi et al., 1999; Wang et al., 2006). In the dominant negative Kuz construct, the Pro-domain and metalloprotease domain are deleted impeding the Notch activation. As per our expectation, we observed slightly higher number of BCs in the egg chambers over expressing the DN KUZ (5.98±0.069 SEM, n=100) over the control samples (4.49±0.072SEM, n=90) (Fig 3K-M). This suggested that Notch activation in the follicle cells negatively affects the BC cell fate specification in developing eggs. After demonstrating that Notch signaling can indeed affect BC fate, we were curious to know if excessive BC fate observed in the Cup mutant could be rescued by restoring Notch activation. To test this, we introduced the gain of function mutation of Notch receptor, *Abruptex ^Ax-16^* (in a heterozygous condition) (Kelley et al. 1987) *in cup^01355^* genetic background and examined if the number of BC are restored to the normal levels. Indeed, it was the case, as we rescued in BC fate specification when heterozygous *N^Ax16^* allele was brought in Cup mutant background (6.616±0.1240 SEM, n=86) (Fig 3N-Q) compared to homozygous Cup mutant (8.551±0.1679 SEM, n=78). This supports our hypothesis that increased BC fate observed in the Cup genetic background is due to suppression of Notch signaling in the follicle cells. Taken together these results above suggest that Cup functions through Notch to modulate the number of AFC that acquire BC fate.

Decrease in levels of NICD and NECD is indicative of inefficient Notch proteolysis(Bland, Kimberly, and Rand 2003; Schroeter, Kisslinger, and Kopan 1998), Hence we next examined the status of Delta ligand to investigate the reason for reduced Notch signaling in the Cup mutant FCs.

### Cup regulates the nurse cell organization and Delta trafficking

One of the primary reasons for examining Delta in the germline nurse cells was that we observed disorganized nurse cell morphology in the *cup^01355^* egg chambers. Unlike the normal round shape nurse nuclei in the WT, we observed elongated, mispositioned nurse cell nuclei in *cup^01355^* egg chambers (Fig 4A-B). Since mispositioned nurse cell nuclei has been reported under conditions of disorganized cytoskeleton(Cooley and Verheyen 2003). we examined the status of actin cytoskeleton in *cup^01355^* mutant egg chambers. We stained the egg chambers with rhodamine-phalloidin to label the F-actin fibers and observed complete absence of distinct actin fibers in the nurse cells of *cup^01355^* egg chambers unlike the control (Fig. 4C-C’). We expressed *cup* RNAi and measured phalloidin intensity in the nurse cells of early-stage egg chambers (< stage 8) to monitor the effect of Cup on the nurse cell cytoskeleton. We observed significantly decreased phalloidin intensity in the Cup depleted early-stage egg chambers as compared to control. This suggested that Cup is important for maintaining the actin cytoskeleton in the early stages of oogenesis too (Fig. S4A-G). We also examined the microtubule network by staining *cup^01355^* egg chambers with a cocktail of α & β -Tubulin antibody and observed that the microtubule framework is also completely disrupted. The tubules were shorter in length and randomly oriented in nurse cell cytoplasm in *cup^01355^* egg chambers (8.82± 0.45 SEM, n=95 fibers of 5 egg chambers) as compared to the control (18.05± 0.47SEM, n=95 fibers of 5 egg chambers) (Fig. 4D-E). Since Cup plays a role in stabilizing mRNAs, we sought to check if actin and α-tubulin mRNA levels are affected in Cup-depleted nurse cells (Broyer et al., 2017). qPCR of total RNA isolated from whole ovaries showed a 1.69 fold and 1.72 fold downregulation in the relative expression of Act5C and αTub84B respectively in *cup^01355^* egg chambers compared to WT (Fig. S4I). We analyzed the distribution of different stages of the egg chamber in WT and *cup^01355^* ovaries by monitoring Cut protein expression, which is dynamic across the different stages of oogenesis (Jackson and Blochlinger, 1997). We did not observe any difference in the proportion of stages of egg chambers isolated from an equal number of WT and *cup^01355^* ovaries. (Fig. S4H). Thus, the difference observed in actin and tubulin mRNA levels is not due to skewed stage distribution in *cup^01355^* ovaries. Together the results above suggest that one of the reasons for the absence of proper nurse cell cytoskeleton in the Cup mutant egg chambers could be the reduced levels of respective transcripts. As cytoskeleton is critical for trafficking of Delta, the ligand for Notch receptor, we focussed our attention on Delta distribution in the germ line nurse cells (Meloty-Kapella et al., 2012.)

**Figure 4:**
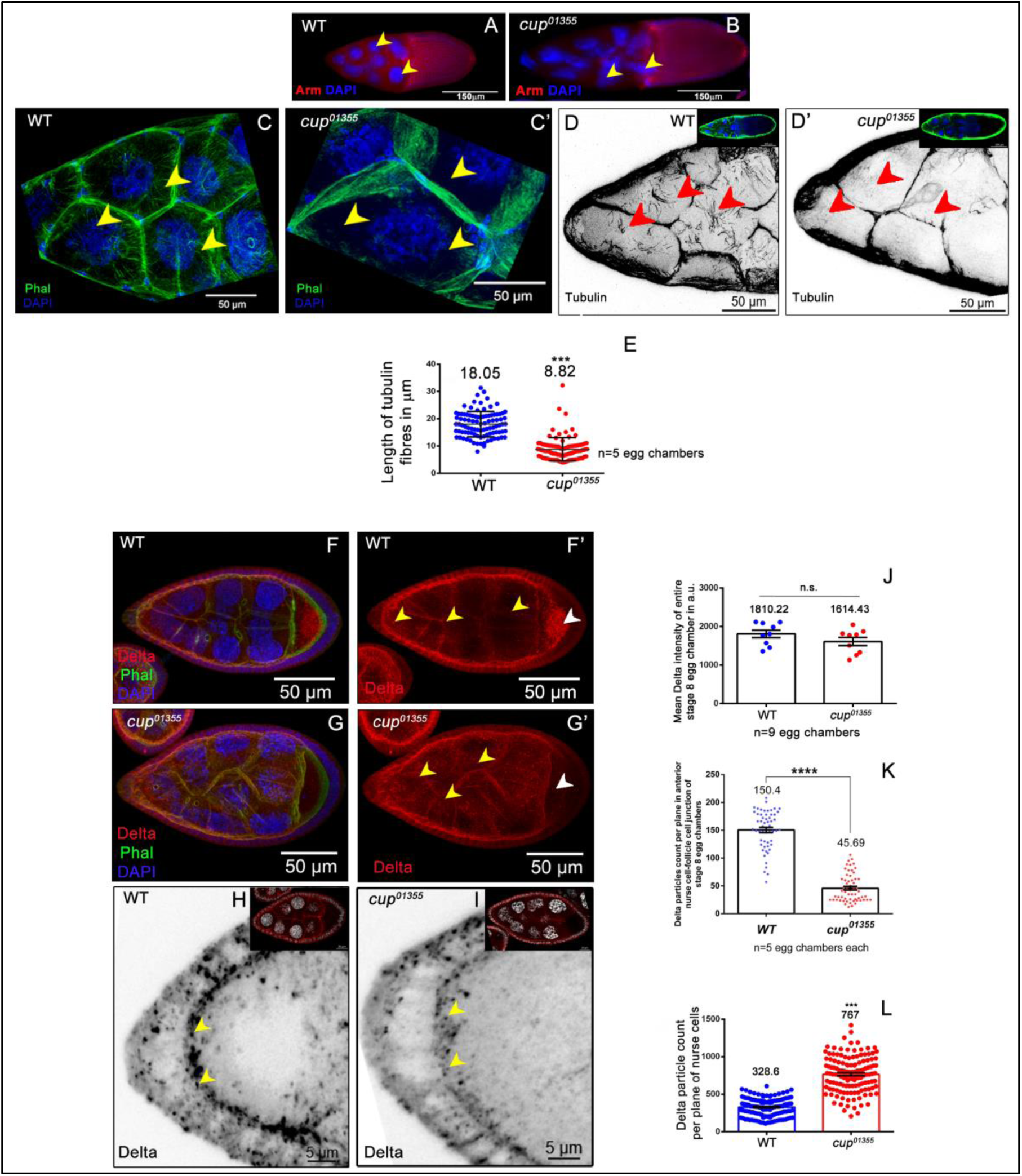
Cup regulates nurse cell organisation which is crucial for Delta internalisation and BC specification. **(A-B)** Nurse cell nuclear morphology is disrupted in *cup^01355^*egg chambers. Nuclei (yellow arrows) are elongated and mispositioned in *cup^01355^* egg chambers compared to round nuclei in wild type, Armadillo (red), DAPI (blue). **(C-C’)** Phalloidin staining of *cup^01355^* egg chambers show absence of distinct actin fibers (yellow arrows) as seen in wild type egg, Phalloidin (green), DAPI (blue). **(D-E)** Tubulin stained *cup^01355^* egg chambers show smaller, randomly distributed tubulin fibres (Red arrows) in nurse cell cytoplasm compared to distinct radially arranged fibers in wild type, tubulin (Black), DAPI (blue). In inset tubulin is in Green. **(F-I)** Delta stained *cup^01355^* egg chambers exhibit more cytoplasmic puncta in nurse cells as compared to wild type (yellow arrows). Oocyte Delta localisation is absent in *cup^01355^* egg chambers as observed in wild type (white arrow). Mean Delta intensity of wild type and *cup^01355^*egg chambers is similar, Delta (red), Phalloidin (green), DAPI (blue). Whereas delta puncta in nurse cell-anterior cell junction is reduced in *cup*^01355^/ *cup*^01355^ over WT.

First, we compared the levels of total Delta protein between WT and *cup^01355^* egg chambers. Unlike our expectation, we did not observe any significant difference in the mean Delta intensity between WT (1810.22±97.63 SEM, n=9 egg chambers) and *cup^01355^* homozygous egg chambers (1614.43±105.29 SEM, n=9 egg chambers) (Fig. 4F-G’,J). Nevertheless, the asymmetrical posterior localization of Delta protein in WT oocyte was absent in the *cup^01355^* mutant egg chambers (Fig 4F-G’). Strikingly, we observed a large number of conspicuous Delta puncta in the cytoplasm of nurse cells of cup01355 mutant egg chambers (767± 20.51 SEM, n=9 egg chambers) over the Wild type egg chambers egg chambers (328.6±10.35 SEM, n=9 egg chambers) (Fig. 4F-G’,L). In addition, we observed very few Delta puncta at the apical interface of AFC and germline nurse cells (control - 150.4± 4.68 SEM, n=5; cup01355-45.69±3.2225 SEM, n=5) (Fig 4H-I,K). Incidentally the anterior most FCs acquire the migratory border cell fate as oogenesis progresses. Delta being a transmembrane protein, it’s enrichment in the cytoplasmic fraction of the Cup mutant egg chambers and absence from apical interface of AFCs, suggested that Delta trafficking is probably perturbed in the Cup mutants.. These results above suggests that Cup mutation affects both the germline cytoskeleton and Delta trafficking in the developing egg chambers.

As proper trafficking of Delta ligand in the nurse cells is critical for Notch activation (López-Schier and St. Johnston 2001), we were curious to examine which part of Delta trafficking pathway was affected in the Cup mutant egg chambers.

### Endocytosis of Delta is critical for proper BC specification

Delta internalization by endocytosis in ligand-producing cells is important for activation of Notch signaling in the receptor producing cells (Langridge and Struhl, 2017; Meloty-Kapella et al., 2012; Okano et al., 2016). As the observed Delta puncta in the nurse cells can be an outcome of defective endocytosis or exocytosis, we first examined the status of these two processes in the Cup mutant nurse cells. This was to investigate which component of the cellular trafficking was defective and resulting in Delta enrichment in the Cup mutant egg chambers. To test this, we carried *ex vivo* live endocytosis uptake assay on egg chambers with an antibody that recognizes the extracellular domain of Delta ligand (c594.4). In the live samples, c594.4 antibody can only bind the extracellular epitope of Delta as it is externally presented while the intracellular Delta fraction remains unlabelled. During chase, the labelled extracellular epitope of Delta is internalized, moves through the endocytotic vesicles along with the ligand. Our rationale was that the enriched Delta puncta observed in the Cup mutant nurse cells will be labelled in the live endocytosis uptake assay if there are defects in endocytosis. While any shortcoming in exocytosis will defer recognition of cytoplasmic Delta by the c594.4 antibody in the Cup mutants in the above assay (Le Borgne and Schweisguth 2003; Giagtzoglou et al. 2012). When we conducted this experiment, we observed a conspicuous apical enrichment and a few randomly distributed cytoplasmic puncta of Delta in the follicle cells of both the wildtype and Cup mutant egg chambers. Strikingly unlike the WT, we observed significantly higher number of cytoplasmic Delta in the nurse cells of Cup mutant egg chambers as observed in fixed sample analysis (Delta particle count: WT-468.3±27.84 SEM n=11, *cup^01355^/ cup^01355^-* 1104±60.88 SEM n=11)(Fig 5A-C). As the cytoplasmic Delta in the Cup mutant nurse cells was conspicuously labelled in the live endocytosis assay, it suggested that defects in Delta trafficking observed in the Cup mutants was predominantly due to impaired endocytosis. To cross check the above observation, we blocked endocytosis *perse* in the germline nurse cells and examined the distribution of Delta ligand. Auxilin (*hsc ^70-4^*) is a J domain protein known to affect Delta endocytosis in the signaling sending cells (Chang et al., 2002; Jia et al., 2015). We down regulated *hsc^70-4^* function in the nurse cells by expressing hsc^70-4^ RNAi with matα-GAL4 driver and observed conspicuous enrichment of Delta puncta in the nurse cell cytoplasm resembling our previous observations with *cup^01355^* mutant egg chambers. This phenotype was also recapitulated when the function of master regulator of endocytosis, Rab5 GTPase (Bucci et al. 1992) was down regulated by RNAi in the nurse cells Fig. 5D-G). These observations support the fact that impaired endocytosis affects Delta trafficking in the nurse cells on similar lines as observed in our live endocytosis assay in Cup mutants.

**Figure 5:**
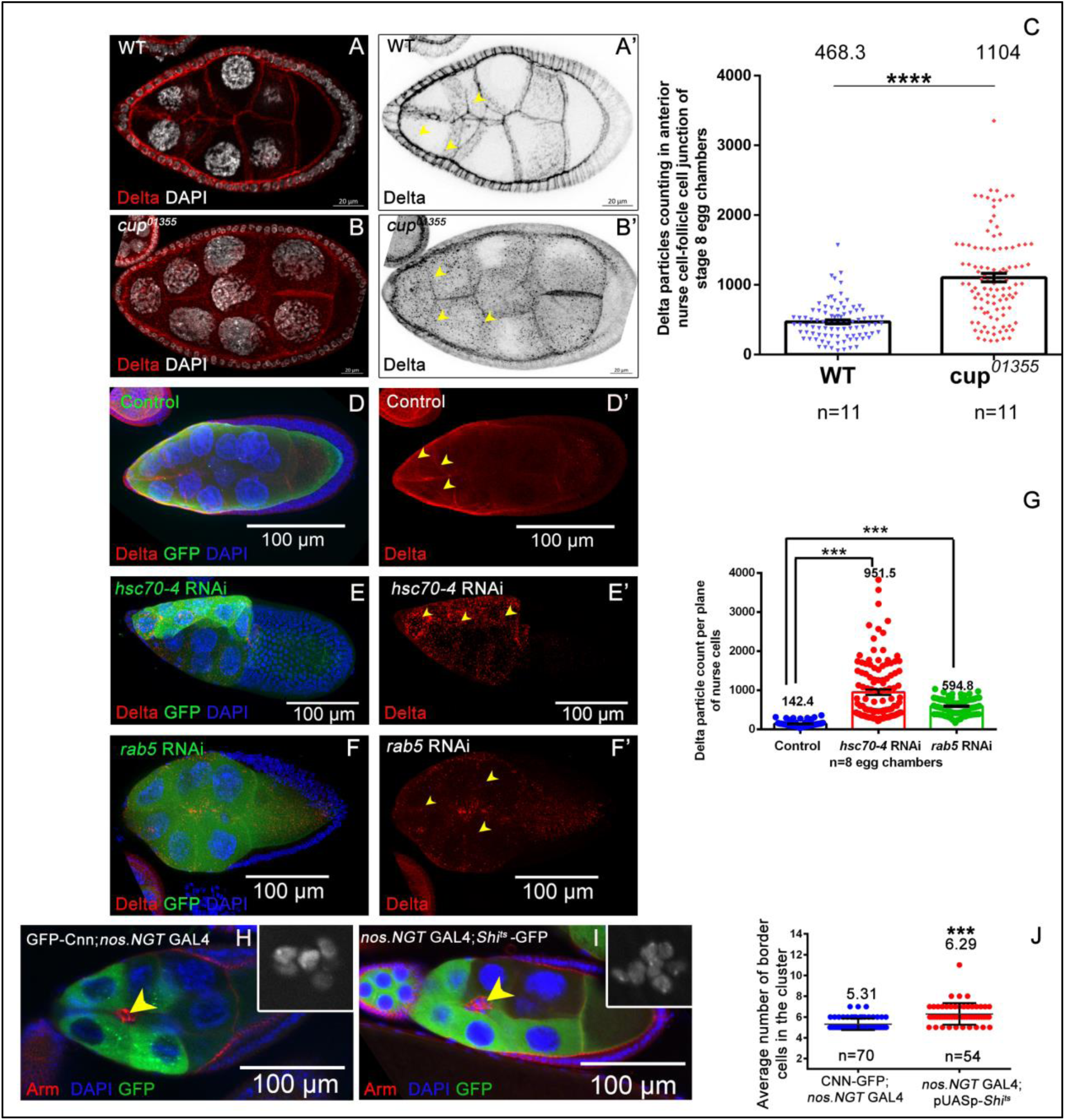
Endocytosis of Delta is critical for proper BC specification. **(A-C)** live Delta internalization assay has shown that *cup^01355^*egg chambers exhibit more cytoplasmic puncta (yellow arrows) in nurse cells as compared to wild type (yellow arrows), Delta (red, black), DAPI (white). **(D-G)** Expression of *hsc70-4* RNAi and *rab5* RNAi in nurse cells using matα-tubulin GAL4-VP16 enriches Delta cytoplasmic puncta (yellow arrows), Delta (red), DAPI (blue), GFP (green). Nurse cells expressing RNAi are indicated by capu.GFP reporter expression. **(H-J)** Downregulation of endocytosis in nurse cells by expressing DN *Shi^ts^*-GFP increases BC number (yellow arrow) compared to control, Armadillo (red), GFP (green), DAPI (blue, inset grey).

Next, we explored the impact of blocking Delta endocytosis on BC fate specification. To do this we down regulated the function of Dynamin, which plays an important role in pinching off vesicles containing Delta-Notch complex in ligand-producing cells (Windler and Bilder, 2010). We expressed a dominant-negative temperature-sensitive allele of Shibire (*Drosophila* homolog of Dynamin) in the nurse cells using the *nos*.NGT GAL4 and assessed the BC fate (Kilman et al., 2009). We used GFP-Cnn as a control reporter to indicate germline expression of *nos*.NGT GAL4. We found that the downregulation of *Shi* activity in the nurse cells increases the number of BCs specified from the AFCs (6.29±0.14 SEM, n=54) compared to the control (5.31±0.06 SEM, n=70) (Fig. 5H-J).

Altogether our results above indicate that endocytosis of Delta ligand in germline nurse cells plays an important role in regulating the number of follicle cells acquiring the BC fate. Any instance wherein Delta internalization is altered, including Cup mutants, causes aberrant BC fate specification. Next, we were curious as to know how Cup function in the germline might affect Delta’s internalization or endocytosis. One possibility could be that lower levels of actin and tubulin may affect the nurse cell cytoskeleton, thus impeding Delta trafficking and Notch activation in the follicle cells. However, over expression of actin and tubulin together or independently failed to rescue the BC fate in the Cup mutant egg chambers (Fig S4J). This suggests that the effect of Cup on the nurse cell cytoskeleton and Delta trafficking may be independent of each other.

Given that endocytic pathway of Delta ligand trafficking is perturbed, we further investigated to identify which component of endocytosis rescues the excessive BC fate observed in Cup mutant egg chambers.

### Rab11 over expression in the Cup germline nurse cells limit BC fate

Endocytosis is a multistep process where the internalized cargo moves to early endosomes, where the cargo is either recycled back to the plasma membrane or directed to late endosome enroute to degradation(Bonifacino and Rojas 2006; Futter et al. 1996; MacDonald, Savage, and Zech 2020; Ullrich et al. 1996). Rab5 GTPase plays a crucial role in biogenesis of endosome and aids in the maturation of early endosomes to late endosomes (Bucci et al. 1992) while Rab11 GTPase facilitates the recycling of the cargo from the early endosomes to the plasma membrane(Dollar et al. 2002; Pasqualato et al. 2004). The cargo that is marked for degradation moves from the late endosome to lysosome with the help of the activity of Rab7GTPase (Guerra and Bucci 2016). As our results suggest that Delta endocytosis machinery is defective, we were curious to know which aspect of endocytosis was impaired in Cup mutants. To investigate this, we overexpressed various constitutively active Rab GTPases (CA) in the nurse cells of the *cup^01355^/ cup^01355^* egg chamber and examined if that could rescue the BC fate specification. The rationale for this experiment was activation of the impaired endocytotic arm in *cup^01355^/ cup^01355^* nurse cells, should be able to rescue the border cells fate in the Cup mutants. We observed that overexpression of Rab11GTPase CA in the *cup^01355^/ cup^01355^* nurse cells rescued number of BCs to the control levels (*cup^01355^*-8.36±0.081SEM, rescue-5.889±0.0.12 SEM, wild type-5.757±0.082SEM, n≥80 egg chambers) (Fig 6A-E, S5A). However, we didn’t observe any significant difference in the BC numbers when Rab5GTPaseCA (8.260±0.1479) and Rab7GTPaseCA (8.913±0.1632) was over expressed in the *cup^01355^/ cup^01355^* nurse cells (Fig 6E; S5D-G). This suggested that activating the recycling component of the endocytosis is able to restore the BC fate to near wild type numbers in the *cup^01355^/ cup^01355^* mutant egg chambers suggesting recycling component of endocytosis in the germline nurse cells is critical for limiting the BC fate from the AFCs.

**Figure 6:**
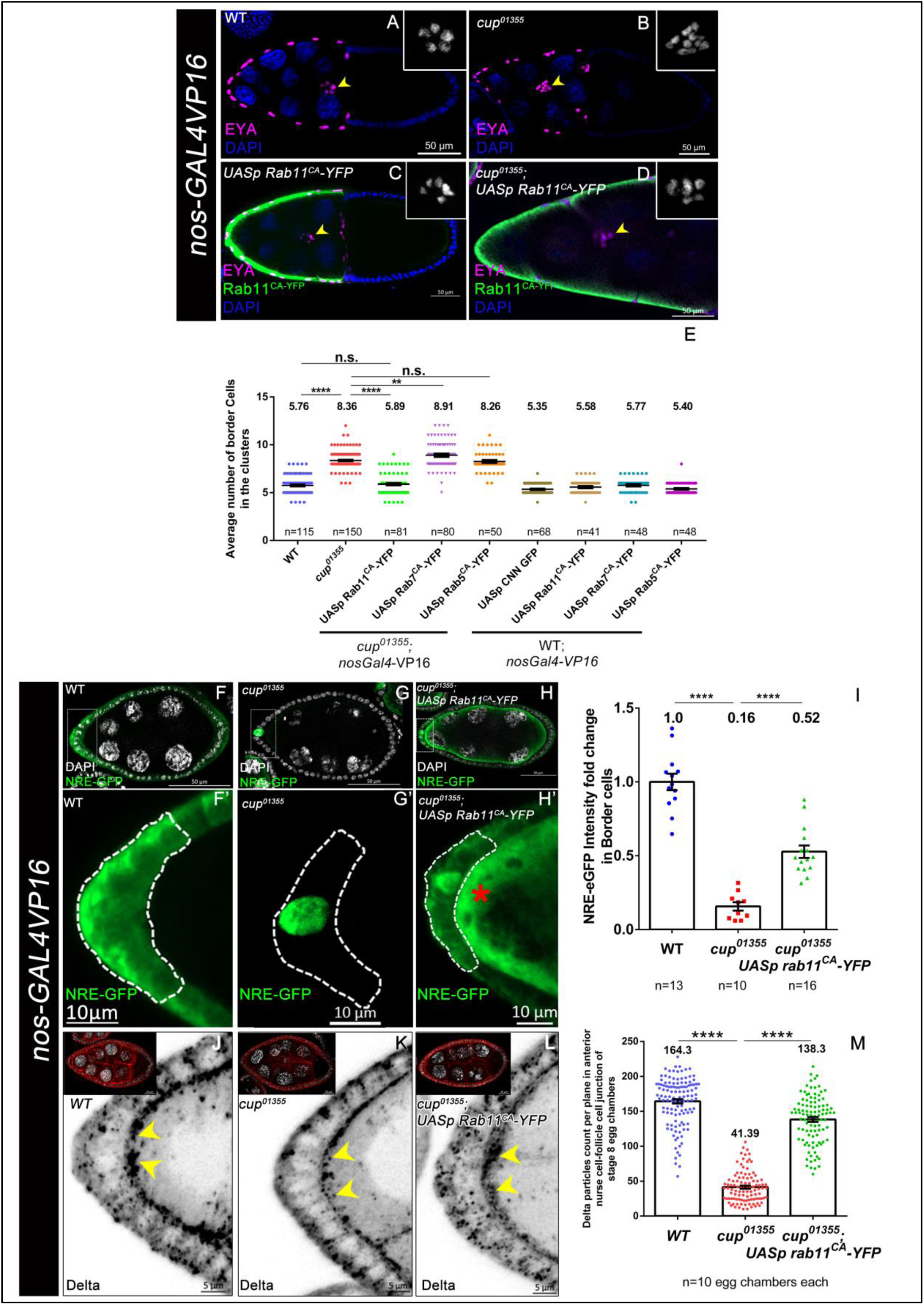
Rab11 overexpression in the cup germline nurse cells limit BC fate. **(A-E)** Stage 10 egg chambers of indicated genotypes stained with EYA in magenta, DAPI in blue, inset grey and YFP in green. yellow arrowheads mark the border cell cluster. No. of border cells are rescued when RAB11CA is overexpressed in nurse cells of *cup01355* egg chambers as compared to *cup01355* egg chambers. **(F-I)** Stage 8 egg chambers of indicated genotypes. White dotted line outlines the AFCs that would be specified as future border BCs. NRE-EGFP expression is in green and DAPI in white. NRE-EGFP intensity fold change is partially rescued when Rab11CA is overexpressed in nurse cells of *cup01355* egg chambers as compared to *cup01355* egg chambers. Red asterisk in nurse cells indicates tagged YFP expression when Rab11CA is over expressed in germ line with germline specific *nos GAL4-VP16*. **(J-M)** Stage 8 egg chambers of indicated genotypes. Yellow arrowheads mark the delta particle localization in anterior nurse cells-follicle cells junction. Delta expression in Red, black and DAPI in white. Number of delta particles in anterior nurse cells-follicle cells junction of stage 8 egg chambers are rescued when Rab11CA is overexpressed in nurse cells of *cup01355* egg chambers as compared to *cup01355* egg chambers.

Finally, we tested if the rescue that we observed in BCs specification, due to the overactivation of Rab11GTPase was indeed due to the restoration of Notch signalling in the AFCs in *cup^01355^* egg chambers? For this, we measured Notch reporter activity by quantifying NRE-EGFP levels and consistent with our expectation, we observed a 0.5 fold upregulation of Notch activity when Rab11GTPase CA was overexpressed in the nurse cells of *cup^01355^* egg chambers as compared to that of control (*cup^01355^* mutant egg chambers) (wild type-1.00±0.05 SEM, n=13, *cup^01355^*-0.16±0.028SEM, n=10, rescue-0.52±0.040 SEM, n=16) (Fig 6F-I). We also observed rescue in the number of Delta puncta at the apical interface of AFC and germline cells of Cup depleted egg chambers that were over expressing Rab11CA (wild type-164.3±3.125 SEM, n=10, cup01355-41.39±2.16SEM, n=10, rescue-138.3±3.396 SEM, n=10) (Fig 6J-M)

Over all our results suggest that stimulating the recycling endocytosis in the nurse cells of the *cup^01355^* egg chambers restores Notch signalling in the AFCs thus limiting JAK-STAT activation and restricting BC cell fate specification.

Altogether, our results suggest that interaction between germline nurse cells and overlying anterior follicle cells regulates the migratory fate of the border cells. Specifically, our data suggests that recycling of Delta ligand in germline nurse cells aids in Notch activation in the anterior follicle cells, thus restricting the domain of JAK-STAT signaling in the AFCs. In Cup mutants, Delta recycling is impeded, thus compromising Notch activation and resulting in excess JAK-STAT signaling and higher number of FCs acquiring migratory BC fate. Altogether, it appears that Cup protein modulates Delta ligand recycling in the germline cells, which aids in non-cell autonomous Notch activation in the AFCs. Once Notch is activated, it restricts JAK-STAT signalling in the FCs, thus optimising the number of cells acquiring BC fate. Our data thus provides a novel insight how the communication between germline and soma may regulate cell fate specification during development.

## Discussion

Cell fate specification is the fundamental basis for generating cellular diversity in developing metazoans. Out of diverse mechanisms, intercellular communication that delineates the migratory individuals in a stationary population plays an important role both in normal development and as well as in various diseased condition including tumour metastasis. Since cellular movement is critical for both normal and pathological events, we employed *Drosophila* oogenesis model to study how intercellular communication between germline and soma results in the specification of BCs from the anterior follicle cells. So far, we know that BC fate acquisition is under strict surveillance of signaling between the somatic FCs. We report a novel finding of how germline-soma interaction limits the size of the migratory BC cluster during *Drosophila* oogenesis. We show that RNA binding protein, Cup, maintains the nurse cell cytoskeleton and regulates Delta ligand trafficking in the germline cells thus facilitating Notch activation in AFCs. In the absence of Cup function, Notch signaling is hampered, leading to elevated STAT and excessive number of AFCs acquiring BC fate (Fig 7)

**Fig 7:**
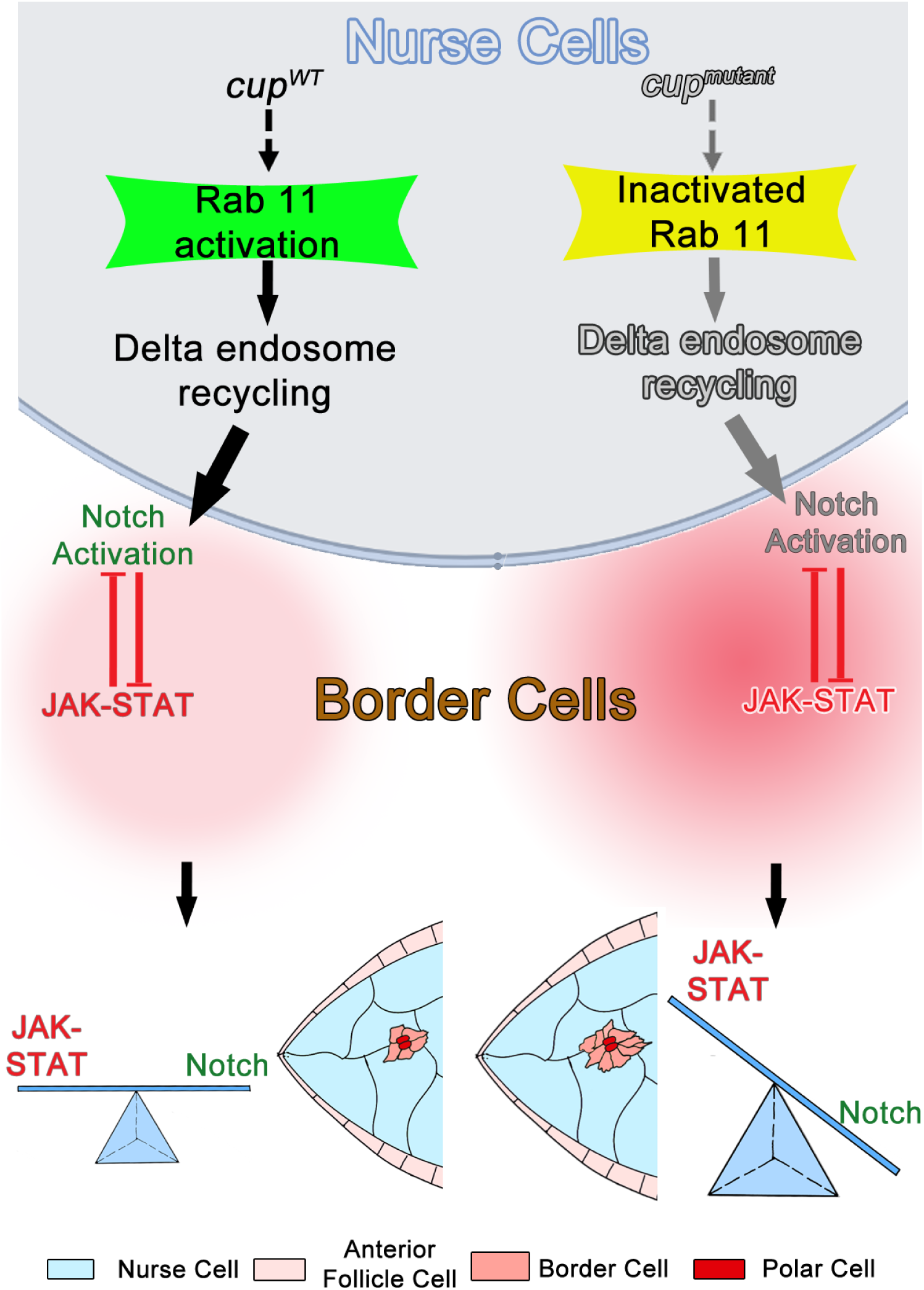
Cup function potentiates Delta recycling in the germline nurse cells. This stimulates non-cell autonomous activation of Notch signaling in the Anterior Follicle cells (AFCs). As Notch and JAK-STAT are antagonistic, a balance between these two signaling cascades aids in transformation of an optimum number of stationary AFCs to migratory border cell fate. In the Cup mutants, Notch signaling is impeded, which results in higher levels of STAT and larger number of AFCs acquiring migratory border cell fate.

.This study was aided by the presence of hypomorphic allele for Cup, which exhibited defect in mid oogenesis. Given that Cup gene plays an essential role in early oogenesis, presence of this allele in particular helped us to bypass the initial steps of Cup requirement and allowed us to examine the role of Cup protein in vitellogenic stages of oogenesis. This won’t have been possible by using classical loss of function mutants of Cup or the Cup RNA interference constructs as both the methods stall the egg chambers in very early stages of development. In the broader context our study highlights the importance of hypomorphic alleles for studying the temporal function of genes that exhibit pleiotropic effect in metazoan development.

Our work highlights how the local environment controls the acquisition of migratory cell fate from a stationary population and gives important insights into the regulation of Notch signaling in vitellogenic egg chambers. Though interaction between the germline and follicle cells have been reported to affect epithelial morphogenesis, this is the first report where germline protein Cup is being shown to non-cell autonomously limit BC fate by restricting Notch signaling in the AFCs. We hypothesize that this interaction probably aids in forming a migrating BC cluster that is just of the right size to efficiently carry the non-motile polar cells to the oocyte boundary for forming the functional micropyle. Besides, our work sheds light upon certain unconventional functions of the germline protein Cup during *Drosophila* oogenesis.

In context of how Cup limits border cell fate in the developing egg chambers? One possibility could be that Cup regulates the germline cytoskeleton which may indirectly affect the Delta trafficking, thus affecting BC fate. However, we don’t believe it be the case, as overexpression of Actin and Tubulin in the Cup mutant nurse cells failed to reduce the excessive BCs observed in Cup mutants egg chambers. Thus, we believe that effect of Cup function on the nurse cells cytoskeleton and on the BC fate are independent of each other. Coupling our results with all the available data on Cup, it appears that Cup performs diverse function in the nurse cells ranging from affecting cytoskeleton stability, regulating the output from maternal mRNAs to modulating Delta trafficking. Thus, it will be worth examining how Cup affects diverse function in the developing germline cells and is there any common over lapping mechanism between these assorted outputs.

Over all we learn that Notch signaling is also required in the mid oogenesis and our data suggests that recycling of Delta is critical for Notch activation. This is significant because there are two proposed models that suggests how Delta trafficking aids in Notch activation in the adjacent cells. The first model suggests that the pulling force generated by Delta – Notch endocytosis in the ligand producing cell facilitates S2 cleavage of Notch receptor, thus activating Notch siganling in the signal receiving cells. While the alternate model proposes that Delta endocytosis coupled with recycling facilitates Delta interaction with Notch receptor thus activating Notch signaling in the receptor producing cells(Bray 2006). In our study, activating recycling component of endocytosis in the germline nurse cells restored Notch signaling and reduced the excessive specification of BCs from the Cup mutant follicle cells. Thus, our data supports the Delta recycling model in the germline nurse cells for activation of Notch signalling in the neighbouring anterior follicle cells during mid oogenesis. Next pertinent question is how Cup is affecting the Rab11GTPase to modulate BC fate from the AFCs? Though we lack complete molecular insight into this aspect, our results allude to that fact that Cup may be regulating the activity of Rab11GTPase rather than it levels *perse*. Our interpretation stems from the fact that over expression of constitutively active Rab11 GTPase exhibits much better effect in limiting excessive BC fate of Cup mutant than overexpression of wildtype copy of Rab11 GTPase (Fig SB-C, H). In future, it may be worth examining how Cup affect Rab11GTPase activity and it will be worth investigating the role of GEFs like Crag in the developing egg chambers.

Notch signaling is an evolutionary conserved pathway in the metazoans, which regulates several aspects of development including cell fate specification, migration, tumor survival by promoting angiogenesis, it would be worth examining if similar modes of germline–soma communication exist in other systems too (Chigurupati et al. 2007; Shi et al. 2005).

## MATERIALS AND METHODS

### *Drosophila* stocks and Crosses

Fly stocks and crosses were maintained at 25 °C and were incubated at 29 °C during GAL4 based experiments. The cup alleles *cup^01355^* (BL-12218), *cup^15^* (BL-29718) were obtained from the Bloomington Stock Centre (BDSC). These two alleles have been characterised by Keyes and Spradling (Keyes & Spradling 1997). Western blot analysis of *cup^01355^* ovaries shows >50% reduction in Cup protein level as compared to wild type (Keyes & Spradling 1997). The *cup^15^* allele is stronger and has been generated by EMS mutagenesis. Western blot analysis of *cup^15^* ovaries shows a very negligible amount of Cup protein as compared to wild type (Keyes & Spradling 1997).

For expression Cup in the germline, pUASp-Cup expression transgenic fly line was generated at the Centre for Cellular And Molecular Platforms (C-CAMP) facility, Bangalore, India. Cup-CDS construct from the BDGP clone LD47924 (Berkeley Drosophila Genome Project), was cloned in the pUASp vector and the construct was used for microinjection. *nos.NGT* GAL4 {Bloomington Stock Center (BDSC 25751)} was used for expressing various transgenes in the germline. The *upd*-lacZ fly stock was a kind gift from Prof. Henry Sun. The stock is generated by inserting a P{lacW} 2851 bp upstream of the 5’ end of upd1 (Tsai and Sun 2004). This construct acts as an enhancer trap reporter enzyme which also harbors a nuclear localisation signal. This expression of lacZ reflects the transcription based on the enhancer activity of the endogenous *upd1* gene. This construct does not reflect the translation status of Upd.

*cup* RNAi (BL-35406), rab5 RNAi (BL-34832), *hsc70-4* RNAi (BL-34836], UAS Kuz DN (BL-6578) Notch response reporter line (BL-30727), N^Ax-16^ (BL-52014), UASp GFP-Cnn (BL-7255), UASp capu.GFP (BL-24763) UASp rab11-YFP (BL-9790), UASp rab11^CA^-YFP (BL-9791), UASp rab5^CA^-YFP (BL-9773), UASp rab7^CA^-YFP (BL-50785) were obtained from BDSC. matα-tubulin GAL4-VP16 (matα-GAL4) was gifted by Daniel St. Johnston. pUASp-Shi^ts^ transgenic line was gifted by Richa Rikhy. For MARCM experiments the stock P{ry[+t7.2]=hsFLP}1, y[1] w[*] P{w[+mC]=UAS-mCD8::GFP.L}Ptp4E[LL4]; P{w[+mC]=tubP-GAL80}LL10 P{ry[+t7.2]=neoFRT}40A; P{w[+mC]=tubP-GAL4}LL7 (BL-42725) was used to cross with *cup^15^* FRT 40A fly stock. BL-5192 was used to recombine FRT40A with *cup^15^* allele. Canton-S flies were used as wild type control flies.

For MARCM experiments 1-2 days old F1 flies were collected and incubated at 37 °C for 1 hour, three times a day with a minimum two-hour interval in between the subsequent heat shocks. Heat shock was given for three consecutive days and flies were fattened at 25 °C after 5 days for 20-22 hrs and then dissected. *cup^15^*homozygous mutant follicle cell clones which spanned the entire anterior end of the egg chamber including the whole border cell cluster was used for quantification of the number of border cells in the cluster. For flip-out experiments, 1-2 days old F1 flies were collected and incubated at 37 °C for 30 minutes three times a day with a minimum two-hour interval. Heat shock was given for three consecutive days, followed by incubation at 25 °C. After 4 days the flies were fattened at 29 °C for 20-22 hours and then dissected.

Clones spanning >50% of the anterior follicle cells with minimum 3 border cell clones were analysed for quantification of number border cells in the cluster.

For GAL4 expression-based experiments, 2-3 days old flies were incubated at 29 °C for 22-24 hours followed by dissection. For mutant-based experiments, 2-3 days old flies were incubated at 25 °C for 22-24 hours followed by dissection.

For downregulation of Shibire, F1 flies bearing *Shi^ts^* expression construct were incubated at 31 °C (non-permissive temperature) for 20 hours followed by incubation at 29 °C for 18hours and then dissected.

For downregulation of Hsc70-4 in nurse cells with matα-GAL4, the flies were fattened at 25 °C for 20 hours followed by incubation at 29 °C for 6-7 hours and then dissected.

### Immunostaining

Ovaries were dissected in Schneider’s media containing 10% FBS (Foetal Bovine Serum, US origin, catalog no. 16000044) and fixed with 4% p-Formaldehyde (Sigma-Aldrich, catalog no. 158127) for 15 minutes at room temperature. Blocking was done with 1X PBS (Sigma-Aldrich, catalog no. P3813) containing 0.3% Triton X-100 (Affymetrix, catalog no. T1001) and 5% BSA (Bovine Serum Albumin, Amresco, catalog no. 0332) for 1hour at room temperature. Mouse anti-Cup antibody was gifted by Prof. Akira Nakamura and used at 1: 10000 dilutions. Rat anti-Slbo antibody was gifted by Pernille Rorth and used at 1:500 dilutions. Rabbit anti-STAT was gifted by Steven Hou and used at 1:750 dilutions. Mouse anti-α-Tubulin antibody (T9026) was obtained from Sigma and used at 1:600 dilutions. Mouse anti-Armadillo (N27A1), mouse anti-Fas III (7G10), mouse anti-Delta (C594.9B), mouse anti-FasII (1D4), mouse anti-EYA (10H6) and mouse β-Gal (40-1a) were obtained from Developmental Studies Hybridoma Bank (DSHB) and used at 1:100, 1:500, 1:200 and 1:100 dilutions respectively. Phosho histone 3 antibody (Cell Signaling Technology, 9713S) was used at 1:150 dilutions. Rabbit anti-GFP (A-11122, Invitrogen) was used at 1:1500 dilutions. Secondary antibodies conjugated with Alexa-488 and Alexa-568 (Molecular Probes) were used at 1:400 dilutions.

For Tubulin cocktail(α+β) staining, Individual egg chambers were dissected in 1X PEM buffer (60mM PIPES, 25mM HEPES, 10mM EGTA, 4mM MgSO4, pH=6.8) and fixed with 10% formaldehyde in presence of 1X BRB80 buffer(Miao, Godt, and Montell 2020) and 1% Tween 20 (Amresco). After fixation wash the sample with 1X PBS containing 1% Triton X-100 and 1X BRB80 buffer for overnight at 4°C. Blocking was done with 1X PBS (Sigma-Aldrich, catalog no. P3813) containing 1% Triton X-100 (Affymetrix, catalog no. T1001) and 5% BSA (Bovine Serum Albumin, Amresco, catalog no. 0332) and 1X BRB80 buffer for 4 hours at room temperature. Mouse anti alpha-tubulin (T9026) was obtained from Sigma and used at 1:800 dilution and anti beta-tubulin (E7) was used at 1:200 dilution incubation in blocking solution for overnight at 4°C. Wash the sample with 1X PBS containing 0.5% Tween 20. Followed by secondary antibodies antibodies conjugated with Alexa-488 and Alexa-568 (Molecular Probes) were used at 1:400 dilutions in blocking solution.

### Measurement of the size of the border cell cluster

For measuring the volume, completely detached border cell clusters at stage 9 that have not reached the oocyte boundary have been considered for analysis.

The border cell clusters are generally spheroid in structure. To measure the size of the cluster, the whole cluster was imaged at 40X magnification taking z stacks at optimal Z intervals suggested by the Zen 2012 software. The image processing and analysis were done using Zen 2012 software. All the stacks were merged together to obtain a 2-dimensional maximum intensity projection (MIP) image. The cluster was outlined in the MIP image, and the maximum and minimum diameter of the cluster was drawn, and the length obtained from the software was noted. The minor and major axes of the spheroid cluster were obtained by dividing the maximum and minimum diameters by 2 respectively. The Border cell cluster volume was obtained using the formula for spheroid (4/3π a^2^b, where a is the major axis, and b is the minor axis. Images were acquired in Zeiss Axio observer 7 with Apotome.2 module.

### STAT intensity quantification

To measure the STAT intensity, the anterior end of stage 8 egg chambers of wild type and *cup^01355^* egg chambers were imaged with center z section passing through the middle of the anterior polar cells. Z sections were captured at regular intervals for both the kinds of samples. The exposure time was kept identical for image acquisition in DAPI (7.2 ms) and Rhodamine channel (4s) (for STAT signal acquisition) for control and experiment samples. All the stacks were merged together to obtain a 2-dimensional maximum intensity projection (MIP) image, and three nuclei on either side of anterior polar cells were outlined, and the mean STAT intensity was noted. Mean STAT intensity was calculated for each egg chamber. The average of Mean intensity for the control and experimental samples was determined and subsequently plotted with statistical tests.

Images were acquired in Zeiss Axio observer 7 with Apotome.2 module and analysed with Zen 2012 software.

### Notch Response Element GFP intensity calculation

For measuring the NRE-GFP levels, the anterior end of stage 8 egg chambers were imaged keeping identical exposure time (GFP channel-400 ms) and other imaging parameters. Stage 8 was identified by depleted levels of Fas2 protein for both wild type and the mutants (Szafranski and Goode 2007). A single follicle cell layer above and below the polar cell containing layer was imaged taking z sections at regular intervals. The z-planes were merged together to obtain a 2D image, and four cells on either side of the polar cell along with the polar cells were outlined as the single region of interest in the anterior end of the egg chamber. The mean GFP intensity of the main body follicle cells (4 cells) was used for background correction. The mean of the corrected GFP intensity for the control and experimental egg chambers was plotted as fold change where we kept control as 1 with statistical tests. Images were acquired in Zeiss Axio observer 7 with Apotome.2 module and analysed with Zen 2012 software.

### Quantification of nuclei in the border cell cluster

The nuclei were labeled using either anti-Slbo antibody or DAPI. The stage 9 or 10 egg chambers, which had detached completely from the anterior end were considered for quantification. The number of border cell nuclei except the two polar cells were counted for every cluster, and the value was plotted. For counting Slbo positive cells, all the nuclei in the cluster, including the polar cells, were counted. In all other experiments, wherever DAPI or EYA was used to evaluate the number of border cells nuclei, the polar cells nuclei were excluded based on their smaller size.

### RNA isolation and qRT-PCR

The RNA isolation was done from ovaries of adult flies using Trizol Reagent followed by cDNA preparation. Status of Cup transcript was evaluated by RT-PCR.

qPCR was done with SYBR Premix Ex Taq from Takara (Catalog: RR420) and StepOnePlus Real-Time PCR System for Act5C and α-tub84B. Primers used are as follows:

Act5C:Forward-5’ACAACGGCTCTGGCATGTG3’, Reverse-5’ GGGACGTCCCACAATCGATG3’,

α-tub84B: Forward-5’CCTTCGTCCACTGGTACGTT3’, Reverse-5’GGCGTGACGCTTAGTACTC3’

Rp49:Forward-5’CTAAGCTGTCGCACAAATGGC3’, Reverse-5’AA CTTCTTGAATCCGGTGGGC3’,

Cup:Forward-5’AATCGTTGGGCCACATCCGA3’, Reverse-5’TCATAGCCAACCGC CTGTGACT3’

Rp49 was used as a house-keeping control.

### Determination of population distribution of different stages of egg chambers

We quantified the proportion distribution of the different stages of egg chambers in the wild type and *cup^01355^* egg chambers in an equal number of ovaries (4 pairs). We stained the egg chambers with an antibody against the protein Cut. Cut is expressed in the follicle cells of previtellogenic egg chambers (up to stage7). During stage 8-10, Cut expression is not detected in the follicle cells and the expression of Cut reappears in the posterior follicle cells after stage10b (Jackson and Blochlinger 1997). We utilised this dynamic expression of Cut to determine the proportion of different stages of egg chambers in wild type and *cup^01355^* egg chambers. We categorised the previtellogenic egg chambers which express Cut in early-stage egg chambers, the stages 8-10 where Cut expression is lost as mid-stage egg chambers and the egg chambers greater than stage10b where Cut expression reappears as late stage egg chambers. The percentage of each of this subset of egg chambers was quantified in both wild type and *cup^01355^* egg chambers obtained from 4 pairs of ovaries, and no significant difference was observed. The experiment was repeated 3 times, and images were acquired in Zeiss Axio observer 7 with Apotome.2 module and analysed with Zen 2012 software.

### Phalloidin intensity calculation

We stained the egg chambers with rhodamine phalloidin and measured the mean phalloidin intensity of nurse cells of early-stage egg chambers (stages 2-7). We imaged the nurse cells of early-stage egg chambers by acquiring Z-sections at regular intervals. All the stacks were compressed and projected as a 2-dimensional maximum intensity projection (MIP) image. The nurse cell region of the egg chamber, excluding the outer follicle cell layer was outlined in the MIP image, and phalloidin intensity was obtained from the software and plotted with statistical tests. Images were acquired in Zeiss Axio observer 7 with Apotome.2 module and analysed with Zen 2012 software.

### Delta puncta quantification

To visualise the Delta distribution, the entire wild type and *cup^01355^* stage 8 egg chambers were imaged taking z sections at regular intervals of 40X magnification in Zeiss LSM 710 confocal microscope or in Zeiss Axio observer 7 with Apotome.2 module. For quantifying the nurse cell cytoplasmic Delta puncta, the z sections encompassing the nurse cells were extracted. The z section images at regular intervals of 0.68 µm were used for counting the puncta to avoid overlapping of puncta amongst the z planes.

The particles were counted using the ImageJ software. The nurse cell area, excluding the follicle cells and oocyte, was outlined for every image. A threshold value was selected on the basis that each Delta puncta was detected as an individual spot, and the background was excluded. The particles were automatically counted by <ANALYZE Particles> option. The average radius of Delta particles was measured and found to be within 0.5 µm. The particle size value range was set from 0.2-1.2 µm^2,^ and circularity was set from 0.5-1. The particles were counted for 9 egg chambers each of control and *cup^01355^* and the total count of Delta particles for each plane was plotted with statistical tests.

### Live delta endocytosis assay

Individual egg chambers were dissected in live imaging media(Prasad et al. 2007). After dissection replace the LIM with mouse anti-delta (1:20 dilution) containing LIM and incubate at 25°C for 1 hour. Wash the sample with LIM for two times and fixed the sample with 4% PFA for 15 mins. Blocking was done with 1X PBS (Sigma-Aldrich, catalogue no. P3813) containing 0.1% Triton X-100 (Affymetrix, catalogue no. T1001) and 5% BSA (Bovine Serum Albumin, Amresco, catalogue no. 0332) for 1.5 hours at room temperature, followed by Secondary antibodies conjugated with Alexa-488 and Alexa-568 (Molecular Probes) were used at 1:400 dilutions.

### Delta intensity quantification

To quantify the total Delta protein, z section images of entire stage 8 control and *cup^01355^* egg chambers (follicle cells and nurse cells) were acquired at regular intervals of 0.43µm in Zeiss LSM 710 confocal microscope. The z planes were merged to obtain a 2D MIP image, and the whole egg chamber was outlined to determine the mean Delta intensity and plotted with statistical tests. Zen 2012 (blue edition) was used to analyse the images.

### *upd*-lacZ intensity quantification

The *upd*-lacZ fly stock was a kind gift from Prof. Henry Sun. *upd*-lacZ consists of a regulatory sequence of Upd gene driving the expression of lacz, which reflects the transcriptional status of Upd locus.

For determining the lacZ protein levels, immunostaining was performed using a primary antibody against the β-Gal protein. The polar cells at the anterior end of egg chambers were imaged at 40X taking z sections at regular intervals keeping equal exposure for experiment and control. All the stacks were merged together to obtain a 2-dimensional maximum intensity projection (MIP) image. The polar cells were outlined in the MIP image, and mean lacZ intensity was obtained using the Zen 2012 software. Images were acquired in Zeiss Axio observer 7 with Apotome.2 module.

### Statistical test

Two-tailed t-test of unequal variance in Excel was used to determine the statistical significance. Standard Error of Mean value was used to plot the error bars. A range used for assigning the p-value is as follows: p value <0.001 is designated as ***, p value <0.01 is designated as **, and 0.05< p <0.01 is designated as *. n= number of egg chambers.

## Acknowledgments

We are thankful to Drs. Henry Sun, Richa Rikhy, Pernille Rorth, Steven Hou, Daniel St. Johnston and Akira Nakamura for providing the crucial fly stocks and antibodies. We thank Bloomington Drosophila Stock Centre (BDSC), Kyoto Stock Center (Japan), Developmental Studies Hybridoma Bank (DSHB), Berkeley *Drosophila* Genome Project and Centre for Cellular And Molecular Platforms (C-CAMP) facility (Bangalore, India) for providing reagents and services. We thank IISER Kolkata imaging facility and in particular Ritabrata Ghosh for help in capturing images in the confocal LSM 710 microscope.

## AUTHOR CONTRIBUTIONS

M.P. conceived the project. B.S., S.A., G.G. and P.D. did the experiments and captured the images. M.P., B.S., S.A. designed the experiments and interpreted the results. B.S. S.A., G.G. and P.D prepared the figures. and the final version of the manuscript.

---

**Figure S1:**
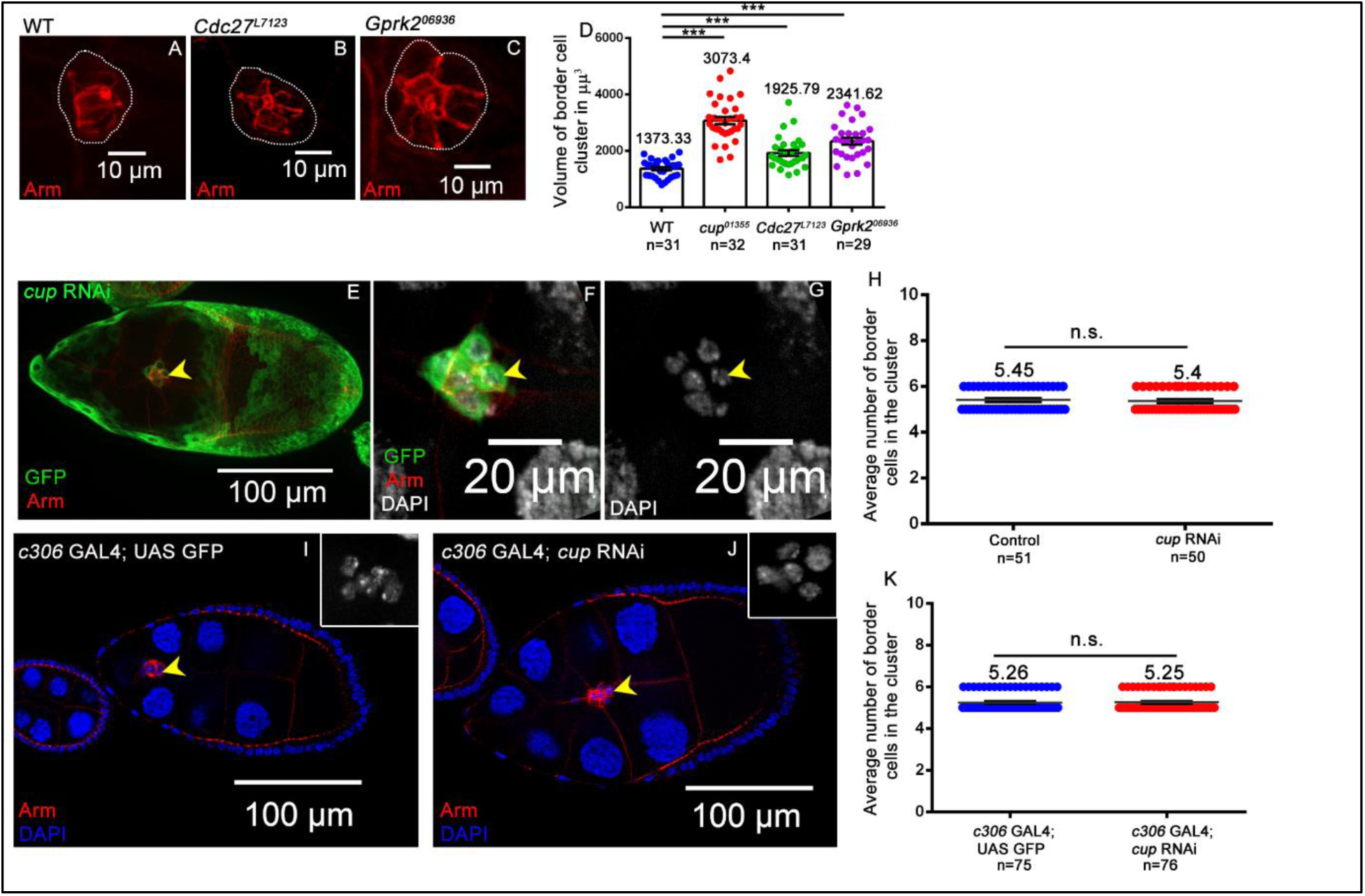
Cup functions in nurse cells to regulate border cell cluster size. (**A-D**) Respective homozygous mutant egg chambers exhibit increased border cell cluster size Armadillo (red), compared to wild type. The white dotted line marks the BC cluster. **(E-H)** Anterior follicle cells along with border cell cluster expressing *cup* RNAi; clones marked by GFP (green), Armadillo (red), and DAPI (blue and grey) does not alter the number of border cells in the cluster compared to egg chambers without clones. The yellow arrow indicates the border cell cluster. **(I-K)** Expression of *cup* RNAi in anterior follicle cells using *c306* GAL4 driver does not alter the number of border cells in the cluster compared to control egg chambers. Yellow arrow indicates the border cell cluster.

**Figure S2:**
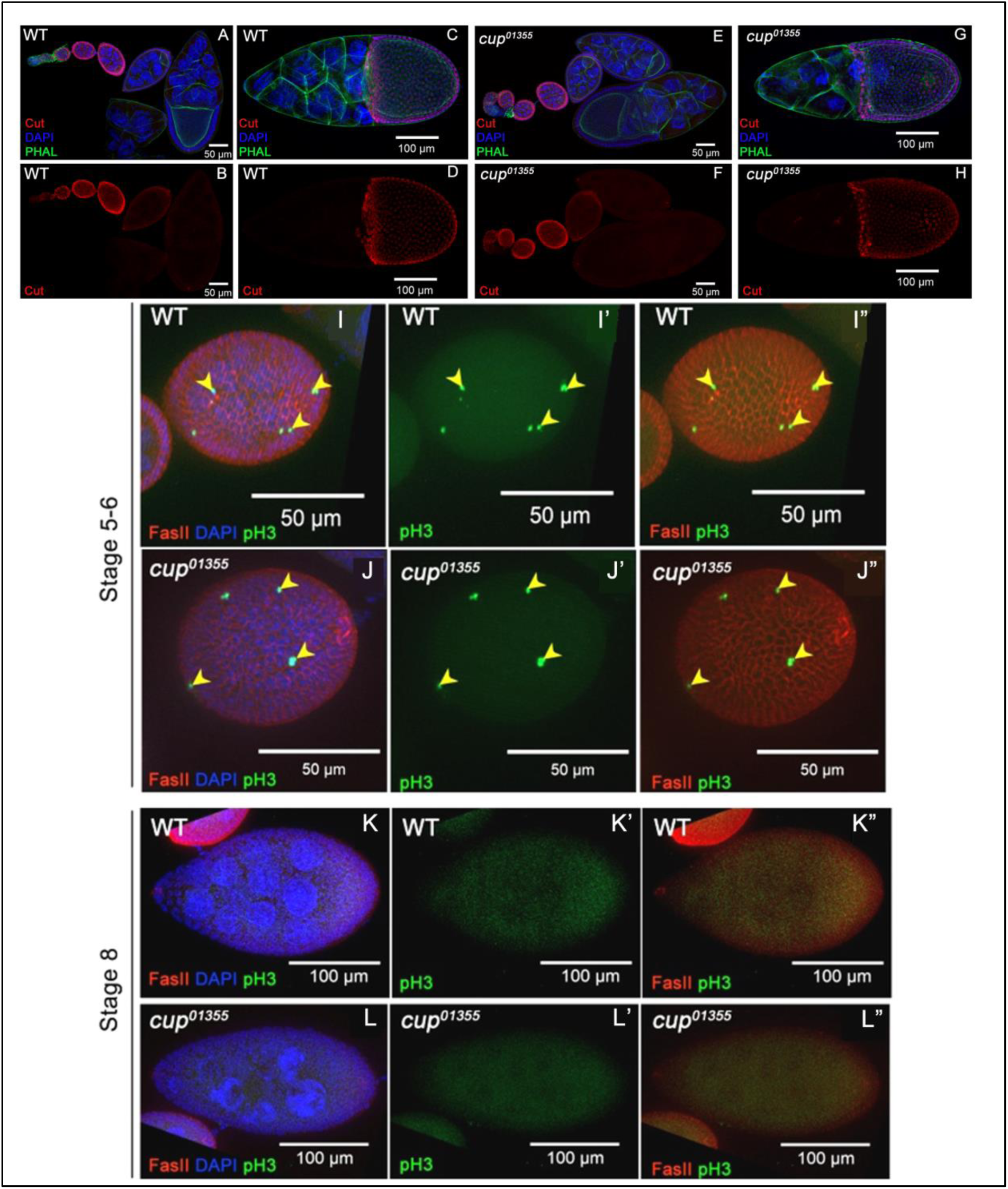
Cup mutation does not alter distribution of different stages of egg chambers. **(A-H)** Expression endoreplication marker Cut in *cup^01355^* early and late-stage (> stage10) egg chambers. Cut (red), Phalloidin (green), DAPI (blue). **(I-L’)** Phospho histone 3 (pH3) staining is observed only in early stage egg chambers (up to stage 6) in both wild type and cup01355 egg chambers (yellow arrow). The presence of FasII indicates it to be early egg chamber. pH3 staining is not observed in stage8/9 egg chambers of wild type and cup01355 egg chambers indicating no cell proliferation after stage 7.

**Figure S3:**
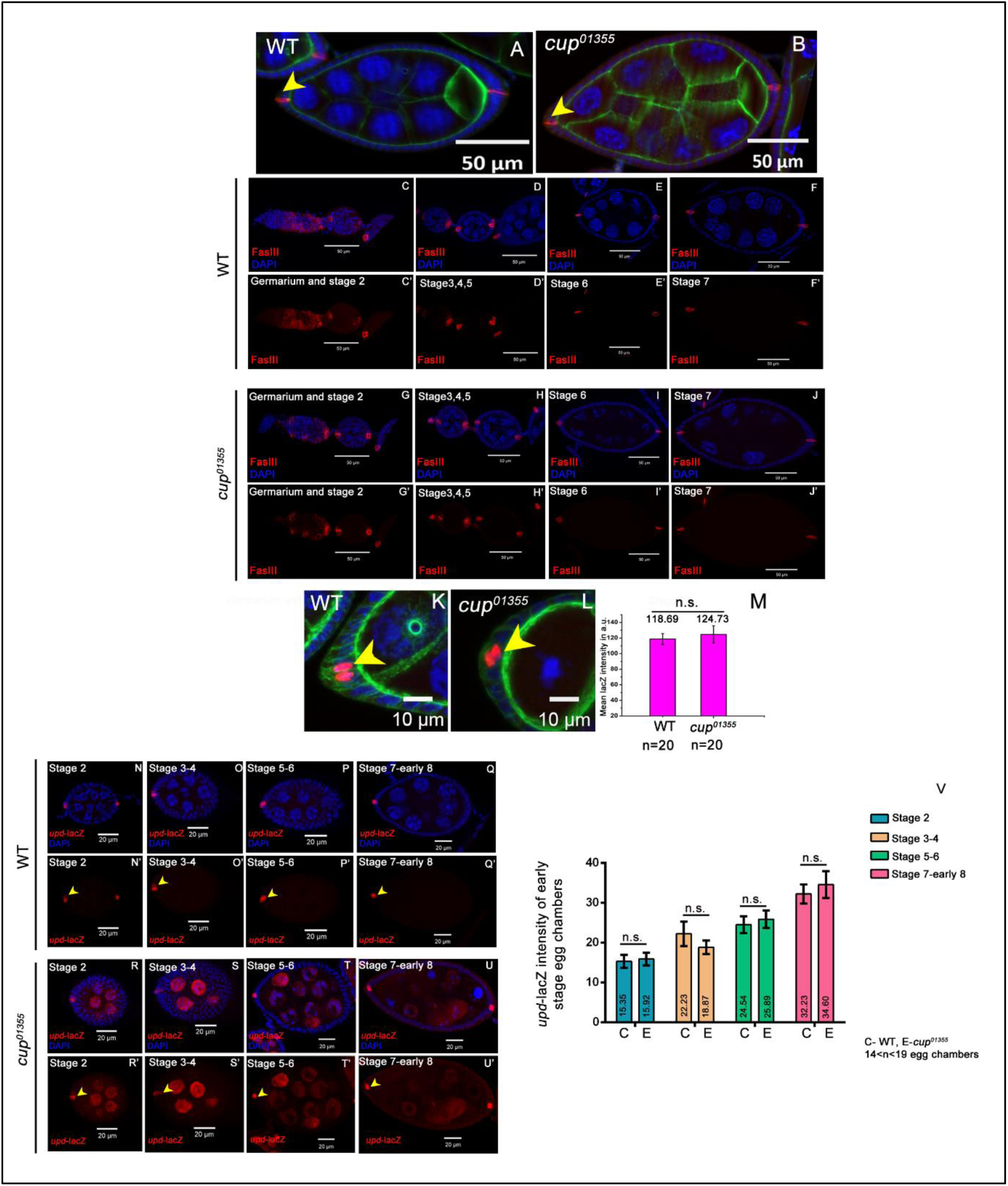
Cup mutation does not affect polar cell specification and Upd ligand production. **(A-B)** Number of polar cells is same in wild type and *cup^01355^*stage 8 egg chamber indicated by FasIII staining (red). Phalloidin (green), DAPI (blue). The yellow arrow indicates polar cells. **(C-J’)** Number of polar cells are same in the early stages of oogenesis (stage2-7) in wild-type and *cup^01355^*egg chambers indicated by FasIII staining (red), and DAPI (blue). **(K-M)** *upd-*lacZ intensity of polar cells is not changed in stage 8 *cup01355*egg chambers as compared to wild type. lacZ (red), Phalloidin (green), DAPI (blue), and yellow arrow indicate polar cells. **(N-V)** *upd-*lacZ intensity of polar cells is not changed in early-stage (2-7) *cup^01355^* egg chambers as compared to wild type. lacZ (red), DAPI (blue), and yellow arrow indicates polar cells. Error bars SEM, t-test.

**Figure S4:**
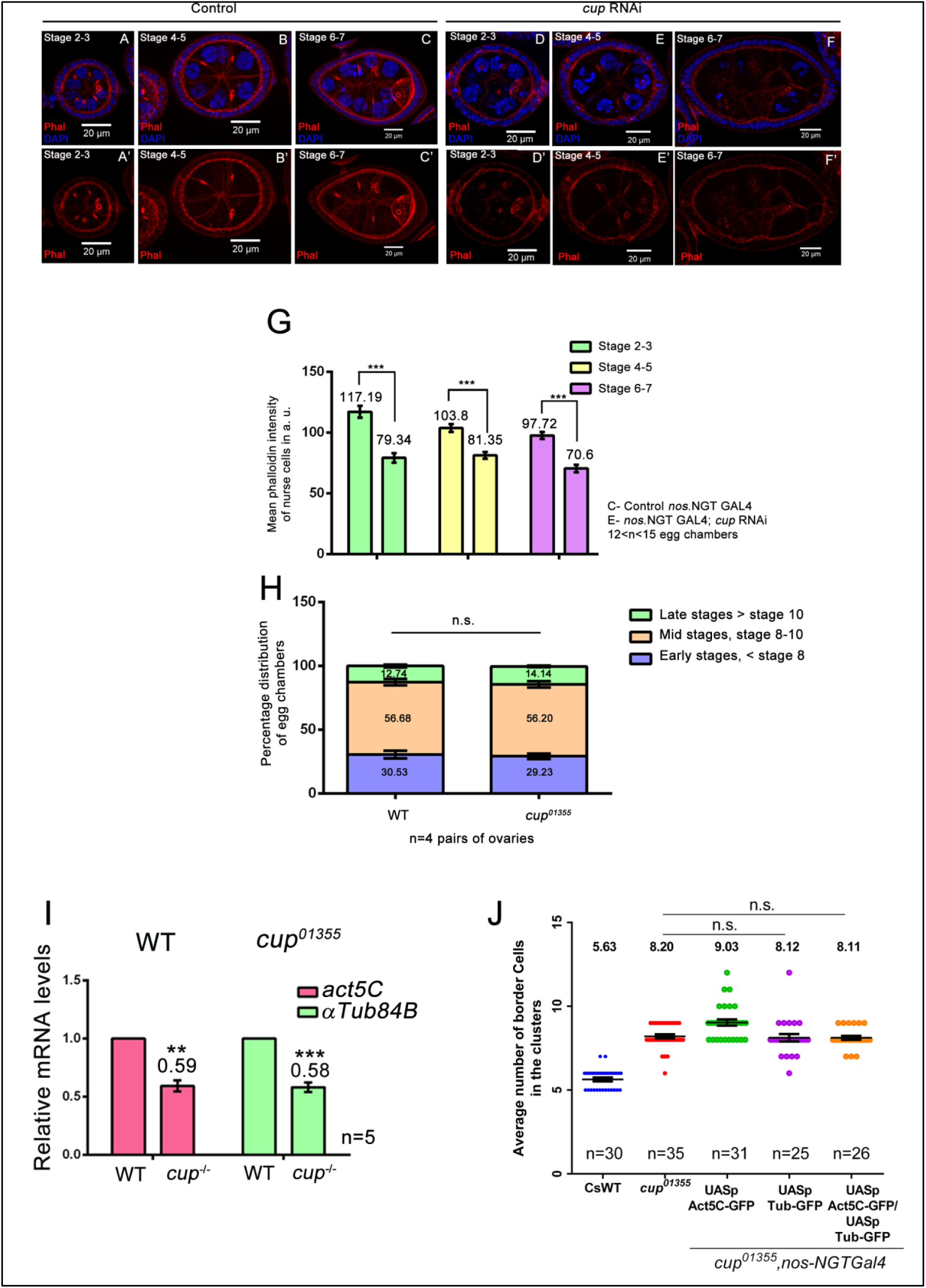
Cup maintains nurse cell cytoskeleton in early stages of oogenesis. **(A-G)** Phalloidin intensity of early-stage egg chambers (2-7) is lowered when cup RNAi is expressed in the nurse cells as compared to control. **(H)** The population of different stages of egg chambers is similar in WT and *cup^01355^*ovaries. **(I)** Graph representing the relative mRNA levels of *Act-5C* and *α-tub84B* in cup mutant and WT egg chambers **(J)** Over expression of actin and tubulin in nurse cells of *cup^01355^* egg, chambers do not rescue border cell number.

**Figure S5:**
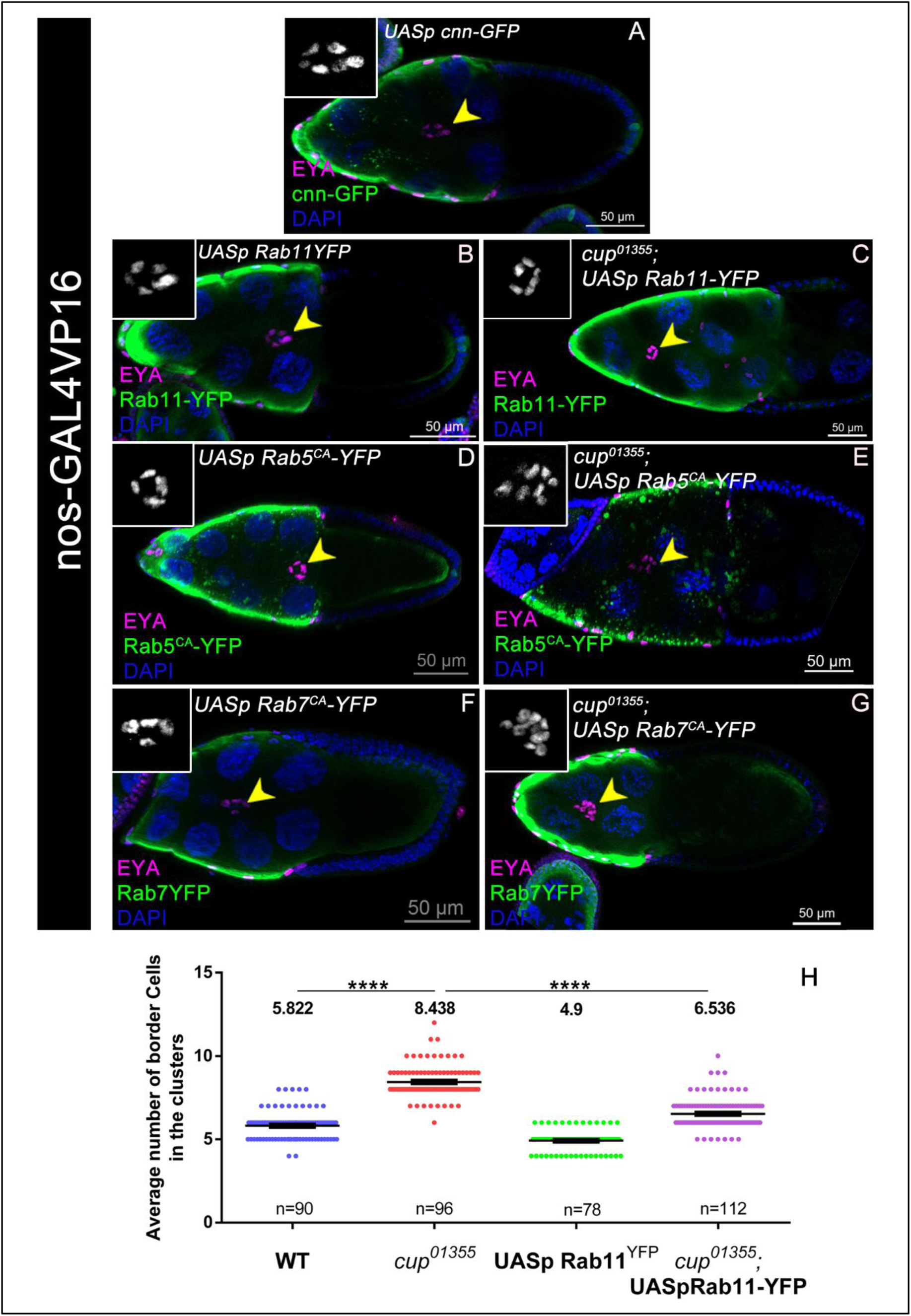
Interaction between Rab GTPases and cup. **(A-G)** Stage 10 egg chambers of indicated genotypes stained with EYA in magenta, DAPI in blue, inset grey, and YFP in green, yellow arrowheads mark the border cell cluster. **(H)** No. of border cells are rescued when WT Rab11 is overexpressed in nurse cells of cup01355 egg chambers as compared to cup01355 egg chambers.

**Table S1:** The list of female sterile lines screened harbouring germline specific gene mutations.

| Germline expressed gene | Allele screened | Homozygous viability |
| --- | --- | --- |
| *Gprk2* | *Gprk2*<sup>06936</sup> | Viable |
| *orb* | *orb*<sup>dec</sup> | Not viable |
| *spn-E* | *spn-E*<sup>hls-03987</sup> | Not viable |
| *pum* | *pum*<sup>01688</sup> | Not viable |
| *E(var)3-9* | *E(var)3-9*<sup>DG08508</sup> | Not viable |
| *bel* | *bel*<sup>neo30</sup> | Not viable |
| *Cdc27* | *Cdc27*<sup>L7123</sup> | Viable |
| *hts* | *hts*<sup>k06121</sup> | Not viable |
| *Ote* | *Ote*<sup>B279</sup> | Not viable |
| *rhi* | *rhi*<sup>02086</sup> | Not viable |
| *psq* | *psq*<sup>KG01598</sup> | Not viable |
| *piwi* | *piwi*<sup>2</sup> | Not viable |
| *Rbp9* | *Rbp9*<sup>P2690</sup> | Not viable |
| *cup* | *cup*<sup>01355</sup> | Viable |

